# Organization of core mitochondrial replication components into multiphasic condensates

**DOI:** 10.64898/2026.03.03.709306

**Authors:** Yuan Yao, Ashish Shyam Tangade, Quentin Livingston, Anupam Mondal, Nidhi Parikh, Jeetain Mittal, Marina Feric

## Abstract

Genomes are organized across several hierarchical levels. Intracellular phase transitions compartmentalize many genomic processes, including transcription and DNA repair. However, little is known regarding how phase transitions contribute to replication, in which long strands of double- and single-stranded DNA need to be coordinated. Here, we investigated the molecular interactions driving the condensation of core mitochondrial replication components into mt-nucleoids. Complex phase behavior emerged among purified mt-replication components: each nucleic acid colocalized with its cognate architectural protein within multiphasic condensates. Using single-molecule experiments, we found that formation of ssDNA increased the partitioning of its cognate protein mtSSB within the condensate, consistent with the preferential localization of mtSSB to replicating mt-nucleoids. To develop mechanistic insights, we built a minimalistic coarse-grained model of mt-replication components that showed how interactions between binary pairs dictate their assembly within condensates. The multiphasic organization of mt-nucleoids has implications for how replication can be spontaneously organized in cells.

**Highlights:**

- Complex co-phase behavior of TFAM-mtDNA depends on their affinity and DNA length
- Mt-replication architectural proteins segregate ssDNA and dsDNA within condensates
- mtSSB selectively partitions into actively replicating nucleoids *in vivo*
- Exposure of ssDNA promotes mtSSB partitioning into TFAM-mtDNA condensates

## Introduction

One defining feature of genomes is their ability to replicate^1^. In doing so, eukaryotic replicative DNA polymerases move at a speed of roughly 1 kb/min with help from helicases that unwind double stranded DNA (dsDNA)^2^. This process thus exposes long stretches (>1 kb) of single strands of DNA (ssDNA), which alone would be susceptible to damage^3–5^. As a counter measure, key architectural proteins coat ssDNA, ensuring the ssDNA maintains its integrity^6,7^. However, little is known how these long genomic strands of dsDNA and the exposed stretches of ssDNA maintain their organization and fidelity with each cell cycle. Replication therefore presents a unique organizational challenge, as dsDNA and ssDNA coexist dynamically, yet interact with distinct binding proteins, raising the possibility that replication-associated assemblies may not be uniform, but instead, internally structured.

Recently, intracellular phase transitions have emerged as a defining organizational principle in genome organization and function, particularly for gene-silenced regions of heterochromatin^8–10^ as well as actively transcribed regions, including the nucleolus^11^ and Pol II mediated transcriptional hubs^12^. More recently, DNA repair machinery has also been described to form condensates around damaged sites^13^. However, how phase separation contributes to the fundamental process of replication and the nature of any potential replication-associated condensates have not been explored.

To study how biomolecular condensates contribute to replication, we turned to a biological system that involves relatively few components: mitochondrial replication. Indeed, mitochondria contain a small, circular genome (mtDNA, 16 kb)^14^ that is packaged primarily by the mitochondrial transcription factor A (TFAM)^15^. The mitochondrial genome is replicated by a two-subunit DNA polymerase (POLG1/2) and a helicase (TWINKLE) that work in tandem^16^, exposing single-stranded DNA that becomes coated by an abundant mitochondrial ssDNA binding protein (mtSSB)^17^. These components assemble into higher-order structures called mt-nucleoids^18,19^, which we found exhibited properties of biomolecular condensates^20^. Mitochondrial replication thus contains each representative biopolymer associated with replication: dsDNA and ssDNA, each with its own architectural protein, TFAM and mtSSB, respectively, providing a minimal system to study how multiple species are organized within replication condensates.

Here, we reconstituted these core mitochondrial architectural components and found that they spontaneously assembled into multiphasic condensates relevant to the apparent organization of mitochondrial replication in cells. With our light microscopy experiments and coarse-grained simulations, we showed that the differential binding affinity of the mt-replication biopolymers resulted in their heterogenous organization within a condensate, in which each nucleic acid associated with its cognate protein within a sub-phase. Specifically, these biopolymers self-organized such that ssDNA was localized internally with mtSSB, surrounded by dsDNA and TFAM. Our *in vivo* experiments indicated that the miscibility of these components was dependent on the activity of the mt-nucleoids. Through our single-molecule DNA experiments, we directly showed how the generation of ssDNA increased the partitioning of mtSSB within a dsDNA-rich condensate. Overall, our results have implications for understanding how replication components assemble within a condensate more broadly.

## Results

### TFAM-DNA affinity determines condensate heterogeneity by locally compacting DNA

To understand the driving forces underlying condensation of replicating mtDNA, we first reconstituted TFAM and mtDNA *in vitro*. TFAM is the core structural protein of mt-nucleoids and is thought to coat the mt-genome largely by binding dsDNA non-specifically^21,22^. In addition, TFAM plays a regulatory role by specifically recognizing the two promoter regions of mtDNA (the heavy strand promoter, HSP, and the light strand promoter, LSP) where it recruits the mitochondrial RNA polymerase (POLRMT) to initiate transcription^23^. These dual binding modes raise the possibility that local differences in TFAM-DNA affinity may influence how mtDNA is packaged within mt-nucleoids. We therefore investigated the dependency of TFAM-mtDNA co-phase behavior on DNA sequence features through an *in vitro* phase separation assay by combining purified TFAM and one of three 100 bp mtDNA fragments: HSP, LSP and non-promoter (NP) sequences (Fig. 1A-C). We visualized the condensates using high-resolution microscopy and found that that promoter sequences increased the nucleation of condensates compared to non-promoter sequences as evidenced by the number of droplets per field of view (Fig. S1A). Overall, the condensates formed with all three DNA fragments were of roughly similar size (Fig. S1B). However, condensates formed with promoter DNA exhibited significantly more heterogenous internal organization as measured by the Pearson correlation coefficient, ρ, between TFAM and DNA fluorescent intensity within the droplet, where lower ρ values reflect an increased extent of spatial demixing between components (Fig. 1D). The higher spatial heterogeneity between TFAM and DNA observed for HSP and LSP condensates (ρ = -0.1 ± 0.4, mean ± s.d., n = 13 droplets and ρ = -0.4 ± 0.2, mean ± s.d., n = 14 droplets, respectively) compared to NP condensates (ρ = 0.3 ± 0.2, mean ± s.d., n = 12 droplets) correlated with the known stronger binding affinity of TFAM for promoter DNA sequences^24^.

**Fig. 1.**
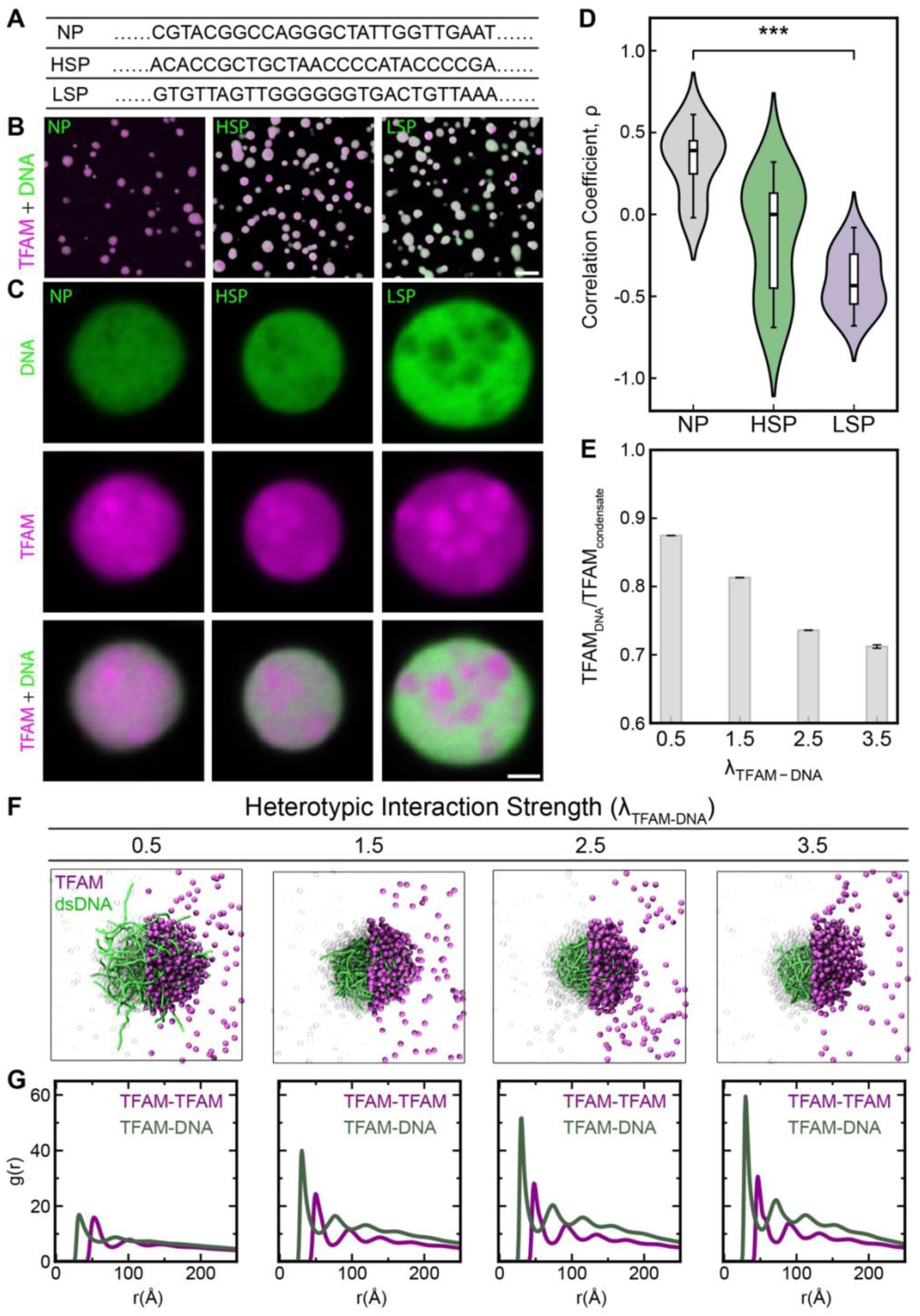
Promoter sequence features potentiate co-phase separation of TFAM and mt-DNA into heterogenous droplets. A) Three 100 bp mtDNA fragments: non-promoter (NP); heavy strand promoter (HSP); light strand promoter (LSP). A ∼20 bp sequence containing promoter and non-promoter sequences is highlighted (see Methods). B) Confocal images of a field of droplets 90 min after mixing TFAM (25 μM) and mtDNA (1.5 uM), with TFAM: DNA ratio at 1 TFAM: 6 bp DNA. Scale bar = 10 μm. C) Confocal images of representative droplets. Scale bar = 2 μm. D) Pearson’s correlation coefficient as a function of mtDNA sequence. ***p < 0.001, t-test. Error bar = S.D. n = 3 independent experimental replicates (each containing n >100 droplets) E) Fraction of TFAM molecules bound to DNA relative to the TFAM in the condensate as a function of heterotypic interaction strength, *λ*_*TFAM*−*DNA*_. F) Representative snapshots from molecular simulations for the condensates of TFAM and 100-mer DNA with increasing strength of heterotypic interactions (*λ*_*TFAM*−*DNA*_). TFAM is represented in violet color, and DNA in green color. TFAM on one semicircular section had been rendered with a transparent setting to highlight the internal organization of DNA with the condensate. G) Radial distribution function *g(r)* for the TFAM-TFAM pairs and TFAM-DNA monomer pairs for the cases of different interaction strength between TFAM-DNA.

Biologically, the heterogenous organization of HSP/LSP-TFAM condensates suggests that promoter regions of mtDNA, which are sites of transcription initiation, may locally alter the internal organization of mt-nucleoids due to their differential TFAM-DNA interactions. Such sequence-dependent effects could provide a mechanism by which specific genomic regions are locally packaged differently within an otherwise non-specific TFAM-DNA condensate. To investigate the physical mechanism underlying the emergence of heterogeneous condensates with increasing binding affinity between TFAM and DNA, we performed molecular dynamics (MD) simulations using a minimalistic coarse-grained (CG) model^25^ to simulate TFAM and 100-mer DNA for condensate formation at the mesoscopic level (see Methods). We varied the heterotypic interaction strength between TFAM and DNA beads (*λ*_*TFAM*−*DNA*_), which acted as a proxy for the effective binding affinity between TFAM and different DNA sequences. In this framework, weaker interactions corresponded to non-promoter DNA, while stronger interactions qualitatively represented promoter DNA, such as HSP and LSP.

To characterize the local mixing within the condensate, we computed the radial distribution function, *g(r)*, for TFAM-TFAM and TFAM-DNA monomer pairs. *g(r)* measures the probability of finding a particle at a certain distance from a reference particle, and its first peak corresponds to the first coordination shell around a particle. As *λ*_*TFAM*−*DNA*_ or the affinity between TFAM-DNA increased, we observed a pronounced increase in the first peak of the TFAM-DNA *g(r)*, correspondingly indicating a higher propensity for TFAM to interact with DNA (Fig. 1F, G). To compare TFAM-DNA heterotypic interactions relative to TFAM-TFAM self-interactions, we computed the ratios of the first peak heights of the TFAM-DNA and TFAM-TFAM *g(r)* (Fig. S1E). This ratio decreased with increasing heterotypic interaction strength, indicating that stronger TFAM-DNA binding promotes spatial heterogeneity within the condensate. We also computed the radial density profile for TFAM and DNA from the center of the condensate to characterize the spatial assembly of these molecules within a condensate (Fig. S1G-J). Lastly, we calculated the ratio of TFAM to DNA density at the center of the condensate (Fig. S1L), which consistently decreased with increasing interaction strength. Together, these measurements from CG simulations support qualitatively similar observations of spatial de-mixing within the condensates as seen in the *in vitro* experiments.

To identify the origin of the heterogeneity that arises with increasing interaction strengths, we examined how TFAM-DNA affinity influences TFAM binding to DNA. With increasing interaction strength of TFAM-DNA, one would anticipate an increase in TFAM bound to DNA; surprisingly, we observed a decrease in the fraction of bound TFAM (Fig. S1K). To investigate this counterintuitive result, we computed the radius of gyration (*R_g_*) of DNA and observed that increasing TFAM-DNA interaction strength led to higher DNA compaction, as indicated by a monotonic decrease in *R_g_* of DNA (Fig. S1F). This increased compaction of DNA reduces the effective surface area of DNA available for additional TFAM molecules to bind, which in turn limits the number of TFAM molecules that can simultaneously bind to DNA, explaining the unexpected trend of decreasing TFAM bound to DNA with increasing interaction strength. Consequently, higher compaction of DNA results in a growing population of unbound TFAM molecules within the condensate with increasing interaction strength (Fig. 1E). These excess TFAM molecules segregated into TFAM-rich regions, giving rise to the heterogeneous internal organization observed in both simulations and experiments. Taken together, these results demonstrate that TFAM-DNA binding affinity is a key determinant of overall condensate architecture, providing a physical mechanism by which variations in the strength of TFAM-DNA interactions can influence the internal organization of mt-nucleoids.

### Long DNA fragments act as scaffolds that globally shape TFAM-DNA condensates

Having established that TFAM-DNA binding affinity gives rise to internal heterogeneity of condensates, we next asked how the length of DNA contributes to condensate morphology and organization. To isolate length effects, we carried out *in vitro* phase separation assay of purified TFAM and mtDNA fragments lacking promoter sequences (NP), thereby removing any contributions of promoter sequence-specific binding effects. DNA lengths ranged from ∼0.1 to ∼10 kb and condensates were imaged using Airyscan high resolution microscopy (Fig. 2A). One prominent observation was that increasing DNA length significantly altered the morphology of the condensates. For example, condensates with shorter DNA (0.1-0.5 kb) appeared mostly spherical (aspect ratio, A.R. = 1), while those with longer DNA possessed more irregular shape in three-dimensional view (Fig. S2D). To better account for the complex morphologies, we quantified the contour of the droplets by measuring the mean negative curvature (MNC) along their perimeter^26^. We observed that MNC increased with DNA length (characteristic length scale, *l*, of ∼15.5 kb), indicating that the condensate adopted increasingly non-spherical morphologies (Fig. 2B).

**Fig. 2.**
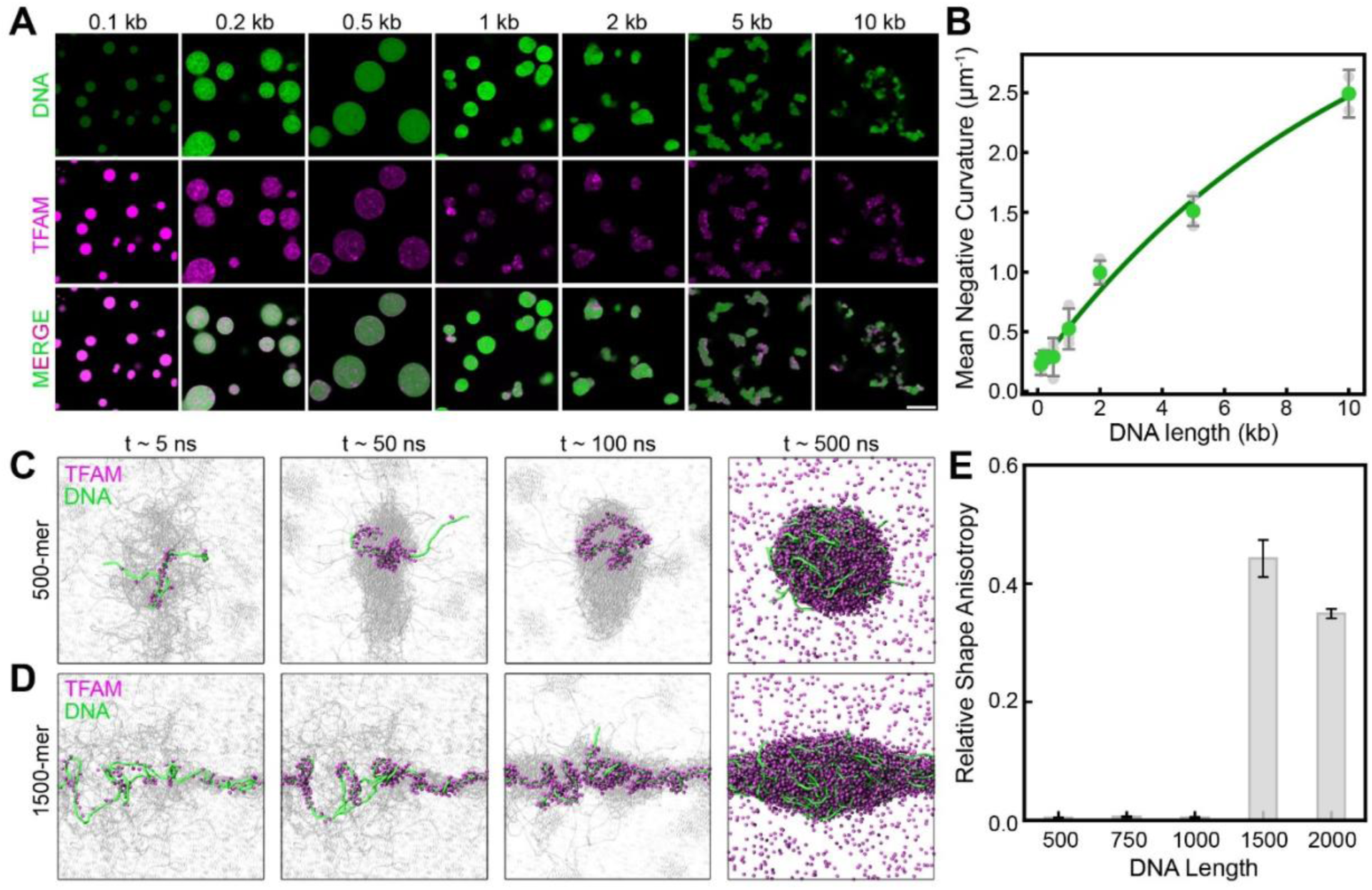
DNA transitions into a scaffold role with increasing length. A) Airyscan images of co-phase separation of 100 ng/uL mtDNA (green, DAPI) of varying length (0.1 to ∼10 kb) and 20 uM TFAM (magenta, STAR RED). The final TFAM: DNA ratio is 1 TFAM : 6 bp DNA. Scale bar = 5 µm. B) Mean Negative Curvature (MNC) of TFAM-mtDNA condensates as a function of DNA length. The data were fit to an exponential curve (green line) corresponding to a characteristic DNA length of ∼15.5 kb. C-D) Formation of biomolecular condensates as observed in the molecular simulations for the case of (C) 500-mer DNA and (D) 1500-mer DNA at a fixed heterotypic interaction strength of *λ*_*TFAM*−*DNA*_ = 0.5. Only a single DNA chain and its bound TFAM molecules are represented in color, whereas other DNA chains and TFAM molecules are represented in grayscale to highlight the mechanism of condensate formation. E) Measure of relative shape anisotropy, *k*, of biomolecular condensates with the increasing DNA length at a fixed heterotypic interaction strength of *λ*_*TFAM*−*DNA*_ = 0.5.

To understand how TFAM and mtDNA assembled into non-spherical condensates, we returned to our minimalistic CG simulation framework. We simulated condensate formation between TFAM and DNA chains of increasing length (500-mer to 2,000-mer), keeping concentrations and affinities constant. Consistent with the experimental system, we used relatively weaker TFAM-DNA interaction strength (*λ*_*TFAM*−*DNA*_ = 0.5) corresponding to non-promoter DNA sequences.

Visualization of the simulation trajectories revealed distinct condensation pathways depending on DNA length. For short DNA (500-mer), TFAM rapidly bound along the DNA and compacted it into a globular structure, which subsequently relaxed into a spherical condensate (Fig. 2C). This behavior closely resembles previously reported condensation mechanism for other DNA-binding proteins such as FUS, where protein-mediated bridging leads to global DNA collapse^27^. In contrast, longer DNA chains (1,500-mer) did not undergo complete global compaction. Instead, TFAM binding induced local compaction of DNA through bridging interactions, while the DNA backbone remained extended at larger length scales. These partially compacted DNA chains acted as elongated scaffolds that recruited additional TFAM molecules, arresting as non-spherical condensates (Fig. 2D). To distinguish the condensate morphologies in our CG simulations, we computed the relative shape anisotropy factor (*κ*^2^), which ranges from zero to one (see Methods), where zero corresponds to a perfectly spherical shape and one corresponds to a rod-like cylindrical shape^9^ (Fig. 2E). Condensates formed with longer DNA exhibited significantly higher *κ*^2^ values, confirming their elongated nature.

We further investigated whether TFAM-DNA interaction strength affects the morphology of the condensates for different DNA lengths. We repeated these simulations at different heterotypic interaction strengths: for all cases, we observed the formation of non-spherical condensates (Fig. S2G), although the minimum DNA length required to observe non-spherical condensate morphology was dependent on the heterotypic interaction strength.

In addition to DNA length-dependent morphologies, condensates formed with longer DNA also exhibited more heterogenous internal organization (Fig. 3A, B). For example, condensates with short DNA fragments (0.5 kb) exhibited speckled organization of DNA and TFAM, in which there were small regions enriched in either TFAM or DNA (*ρ* = 0.0 ± 0.1, mean ± s.d, n = 71 droplets). However, with increasing DNA lengths, such as 5 kb, we found TFAM to be highly variable in the droplets, corresponding to negative correlation coefficients, ρ (*ρ* = -0.2 ± 0.2, mean ± s.d, n = 82 droplets), yielding a characteristic length of *l* = 0.2 kb.

**Fig. 3.**
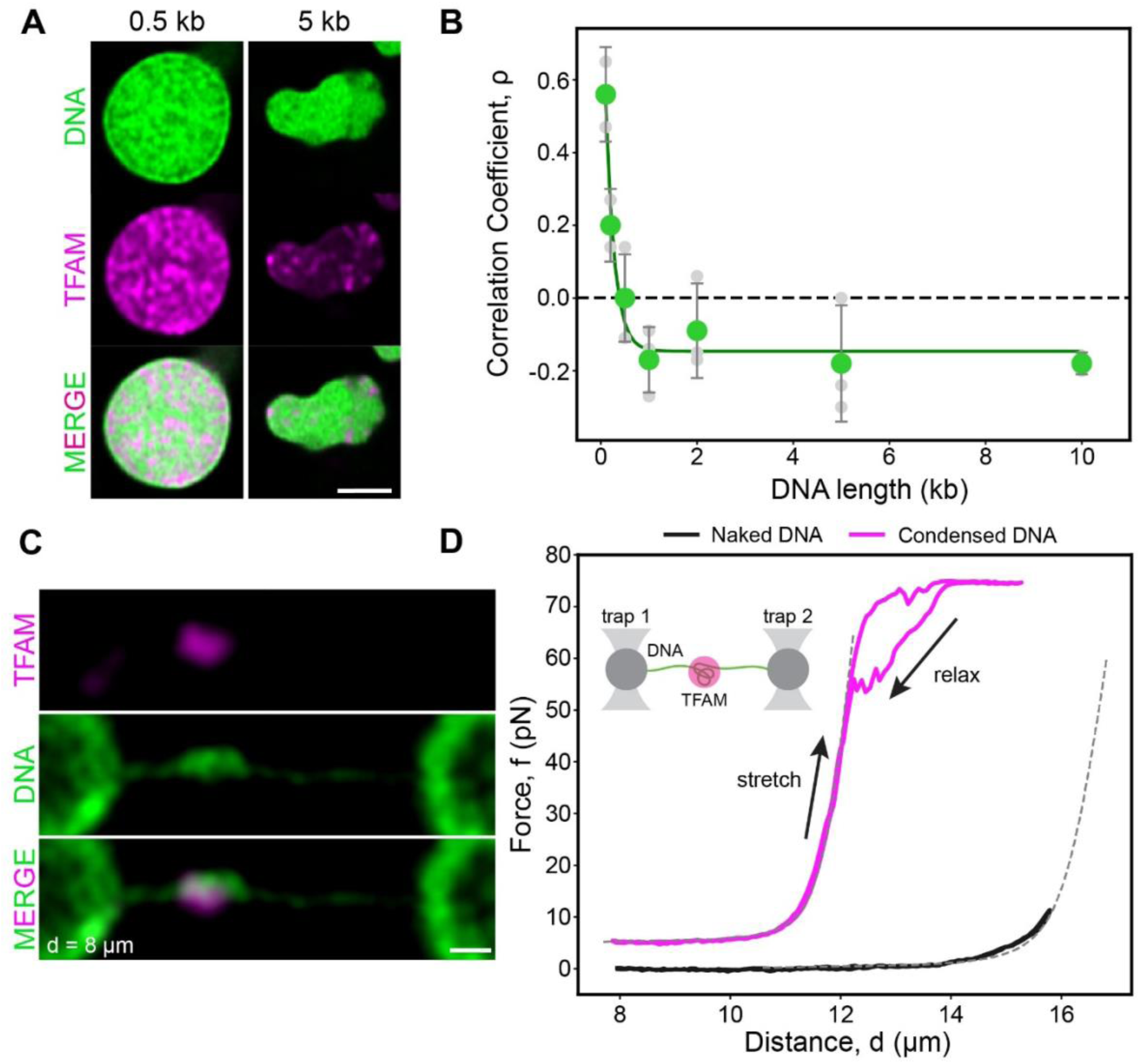
Surface forces needed to compact DNA into a TFAM-rich condensate. A) Airyscan images of representative TFAM-mtDNA droplet with 0.5 kb and 5 kb DNA. Scale bar = 1 µm. B) Pearson correlation coefficient between TFAM and mtDNA as a function of DNA length. The data were fit to an exponential curve (green line) corresponding to a characteristic DNA length of ∼0.2 kb. C) Confocal microscopy image of TFAM (magenta) condensate compacting *λ*-phage DNA (green) tethered between to micron-sized beads (left, right). End-to-end distance, *d*, = 8 µm. Scale bar = 1 µm. D) Force extension curve TFAM-bound DNA (magenta) and naked DNA (black). Dotted line is worm-like chain (WLC) model for non-condensed DNA (L= 16 µm for naked DNA, L = 12 µm for condensed DNA). Inset is schematic diagram of optical tweezer design.

Together, these measurements indicate that DNA length not only determines condensate shape but also modulates internal organization, acting as a second independent parameter alongside TFAM-DNA affinity. While TFAM-DNA affinity determines how TFAM locally packages DNA, DNA length sets a global constraint on how the genome can be compacted. Notably, dependence of MNC of condensate with DNA length shows a characteristic length (*l* ≈ 15.5 kb, Fig. 2B) which is comparable to the size of the full mt-genome (∼16 kb), suggesting that mtDNA is intrinsically long enough to act as a structural scaffold rather than a compact, globular object within mt-nucleoids.

### TFAM condensates exert high forces to compact DNA

To quantify the forces driving TFAM-mediated compaction of DNA into a heterogenous condensate, we performed single-molecule experiments with λ-phage DNA (∼50 kb) and TFAM using dual optical traps. We tethered a λ-phage DNA molecule to two micron-sized beads, and using a microfluidic device, we formed a TFAM condensate on the tethered DNA (see Methods). Using confocal imaging, we observed a high DNA intensity (ρ = 0.7 ± 0.3, mean ± s.d., n = 5 droplets) within the TFAM condensate (Fig. 3C), indicating that DNA was indeed sequestered into the TFAM condensate.

To determine the force required for extension of the condensed λ-phage DNA, we next performed several stretch-relaxation cycles by moving the optical tweezers relative to each other. As a control, we computed the forces required to extend the λ-DNA in the absence of TFAM, referred to as ‘naked DNA’ (Fig. 3D). We found forces larger than over an order of magnitude were needed to pull DNA compacted by TFAM condensates (F = 67 ± 14 pN, mean ± s.d., n=7 droplets) when compared to naked DNA (F = 5 ± 1 pN, mean ± s.d, n = 5 DNA strands) for a given extension (*d* = 15 *μ*m) (p-value = 2 × 10^−6^) (Fig. 3D).

To understand the mechanism driving the significantly higher forces required to extend the compacted DNA within the condensate, we first fit the experimentally determined force-extension curve of DNA to the worm-like chain (WLC) model using the contour length (*L_c_*) of DNA as the fitting parameter^28^ (see Methods). First, we applied the WLC model to naked DNA: the resulting *L_c_* required to fit the naked DNA force-extension curve was ∼16 µm, which was consistent with the known contour length of 50-kb λ-phage DNA^26,27^. In contrast, the force-extension response of condensed DNA was accurately described by using a significantly shorter *L_c_* of ∼12 µm (Fig. 3D), reflecting the length of uncondensed DNA. Thus, the higher forces observed in the presence of a bound condensate were consistent with the resulting reduction in length of exposed DNA: since there was a fraction of DNA that was packaged in the condensate, the remaining uncondensed DNA led to effectively shorter lengths of DNA that were available for stretching.

Notably, at higher extensions (*d* ≥ 12 µm), we observed that the force plateaued at ∼75 pN (Fig. 3D), while the uncondensed length of DNA was fully extended at this distance. This high-force plateau in the presence of the condensate indicates that the condensed DNA can withstand much higher forces. Together, the strong agreement of the WLC when adjusted for DNA length with our force-extension curve and the plateauing of force upon extension of the condensed DNA support the idea that surface and bridging forces applied by the condensate prevent the enveloped DNA from extending upon applied force. In support, we observed that the fluorescent signal of DNA remained stable within the condensates even upon stretching (Fig. S3E, F, Supplementary Video S1), suggesting the DNA continued to be packaged within the condensate, withstanding the applied force. Together, our single-molecule experiments on TFAM-DNA condensates suggest that their heterotypic interactions result in high forces, ensuring stable compaction of the mt-genome within the mt-nucleoid.

### Core components of mitochondrial replication co-phase separate to form multiphasic condensates

We next sought to understand how these properties contribute to the spatial organization of mtDNA replication. To do so, we purified the core mt-replication architectural biopolymers involved in early replication events: the ssDNA-binding protein mtSSB and ssDNA. We found that mtSSB formed prominent condensates both on its own and with ssDNA (Fig. S4A-C), indicating that mtSSB has an intrinsic ability to drive phase separation. We then combined them with our established TFAM/dsDNA *in vitro* condensates to create a four component system (see Methods). We note that for these experiments, our dsDNA and ssDNA templates included the *CYTB* gene (∼1 kb) to capture the early replication events from the OriH^29,30^.

To systematically dissect interactions within this system, we first performed *in vitro* phase separation experiments on all binary combinations of the four-components: TFAM, mtSSB, dsDNA, and ssDNA. Excluding the ssDNA-dsDNA pair, all binary mixtures formed condensates, yet with different extents of miscibility (Fig. 4A). Among the five binary mixtures, mtSSB and its cognate ssDNA formed the most well-mixed condensates (*ρ* = 0.7 ± 0.2, mean ± s.d., n = 105 droplets) followed by mtSSB-dsDNA mixtures (*ρ* = 0.3 ± 0.3, mean ± s.d., n = 65 droplets) (Fig. 4B). Notably, TFAM and mtSSB co-localized into the same condensate, yet exhibited heterogeneous internal organization (*ρ* = -0.1 ± 0.2, mean ± s.d., n = 93 droplets), in which the two proteins occupied distinct regions within the same droplet, consistent with multiphase behavior^20^. Importantly, this observation shows that even in the absence of DNA, the core architectural proteins TFAM and mtSSB can drive multiphase organization. Such protein-protein driven de-mixing suggests that structural features of the mt-nucleoid are likely encoded at the level of protein interactions, before the addition of DNA.

**Fig. 4.**
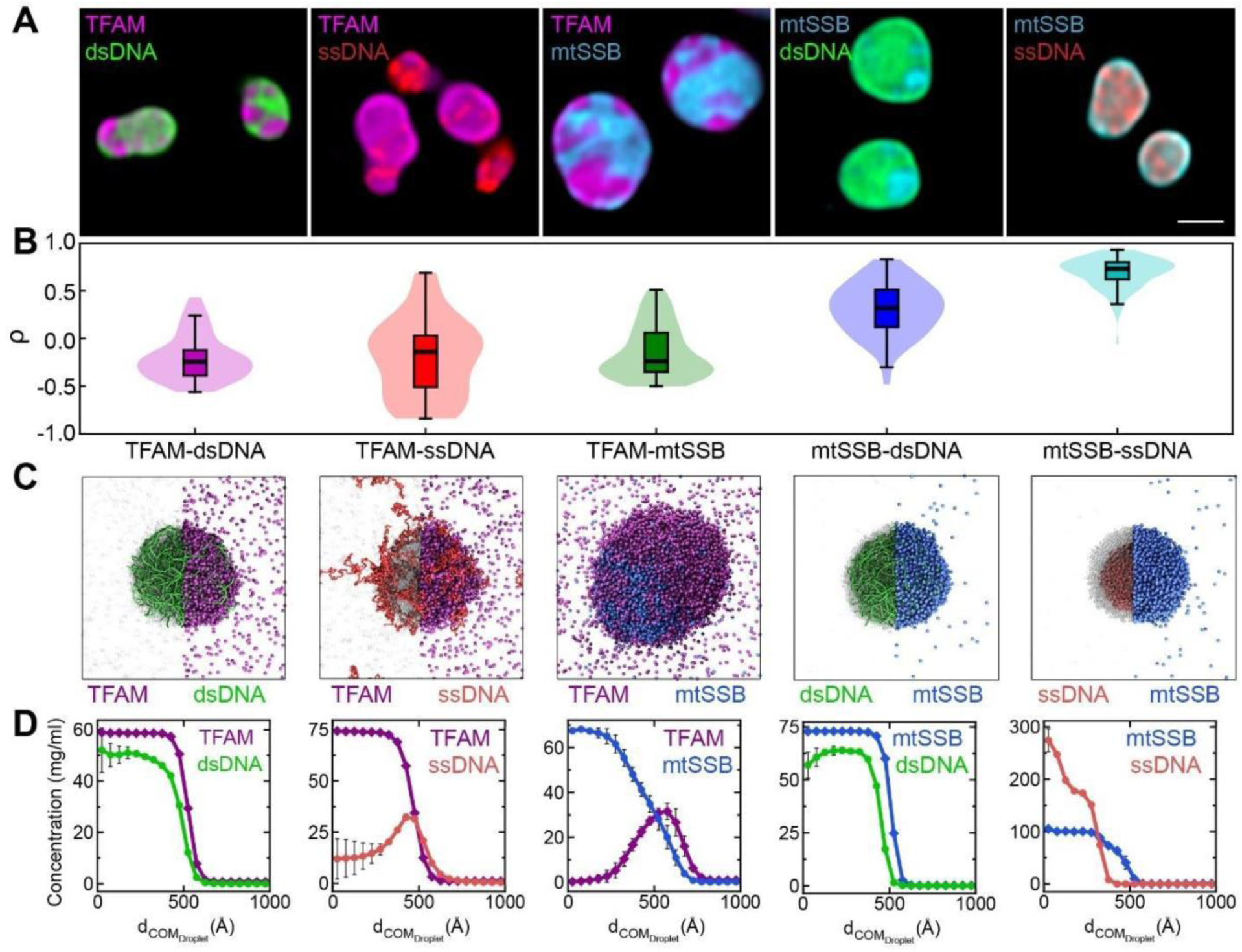
Binary mixtures core mt-replication components display diverse phase behavior. A) Airyscan images of *in vitro* droplets made from binary mixtures of mt-replication components: TFAM-dsDNA, TFAM-ssDNA, TFAM-mtSSB, mtSSB-dsDNA, mtSSB-ssDNA, ssDNA-dsDNA. 20 µM TFAM (magenta, STAR RED), 20 µM mtSSB (cyan, Alexa Fluor 488), 20 nM ssDNA (red, Alexa Fluor 594), and 20 nM dsDNA (green, DAPI). Scale bar = 1 µm. B) Pearson correlation coefficient between components of each binary mixture in (A). C) Representative snapshots from simulations for the binary mixture simulations, indicating the assembly of the two components within the condensate. Protein molecules in half of the condensate are represented as transparent to showcase the internal arrangement of DNA. D) Radial density profile of each component in the simulations of binary mixture shows the localization of the components across the condensate from the center of the condensate.

To model the composition of mt-replication, we then combined all four abundant architectural components to yield a quaternary system of purified TFAM and mtSSB with ds- and ssDNA fragments. Consistent with our model of the mt-nucleoid as a distinct phase, all four components self-assembled into droplets that exhibited distinct heterogenous internal organization, with significant variability in their relative degrees of miscibility (Fig. 5A). The most striking observation was that dsDNA preferentially exhibited peripheral localization, while ssDNA was encapsulated, mostly within the droplet interior. Specifically, ssDNA still highly colocalized with its cognate binding protein mtSSB in the interior, but segregated from dsDNA (*ρ* = -0.2 ± 0.2, mean ± s.d., n = 55 droplets). Consistently, TFAM appeared to colocalize with dsDNA to a similar extent as in the binary droplets (*ρ* = -0.3 ± 0.2, mean ± s.d., n = 55 droplets). When considering the proteins relative to each other, we found that TFAM and mtSSB exhibited partially increased miscibility in the quaternary system compared to their binary mixture, as evidenced by the increased correlation (from ρ = -0.1 ± 0.2 to ρ = 0.2 ± 0.2, mean ± s.d. n = 55 droplets) (Fig. 5B). Together, these changes in co-localization suggest that the presence of ds- and/or ss-DNA partially stabilizes interactions between the two proteins, reducing their tendency to de-mix.

**Fig. 5.**
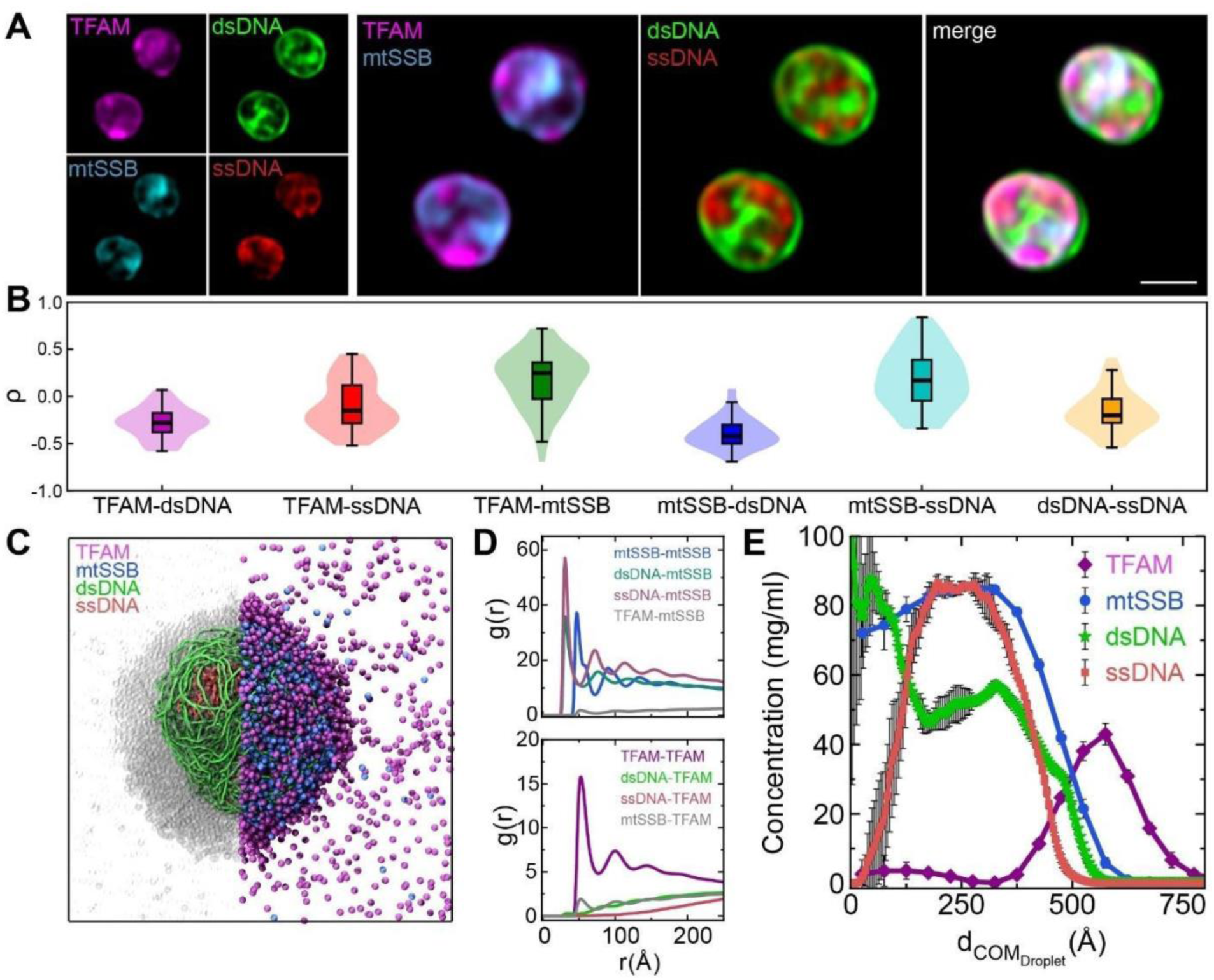
Core mt-replication components spontaneously self-organize into multiphasic condensates *in vitro*. A) Airyscan images of *in vitro* droplets containing all four core mt-nucleoid components: TFAM (magenta), mtSSB (cyan), ssDNA (red), and dsDNA (green). Scale bar = 1 µm. B) Pearson correlation coefficient between each pair of components in four-component droplets. C) A representative snapshot from the simulation of a quaternary mixture of TFAM, mtSSB, ssDNA, and dsDNA. D) Radial distribution function, *g(r)*, for the pair of components with the simulation of the quaternary mixture. The internal organization of the condensate is represented by making proteins transparent in one hemisphere of the condensate. E) Radial density profile of each component in the quaternary mixture from simulations.

To gain mechanistic insight into how effective interactions among the four components drive this organization, we used our minimalistic CG model. We parameterized the effective pairwise interactions in this model by using the observations from *in vitro* experiments of binary mixtures by regenerating qualitatively identical organization of components as observed in experiments through simulations (Fig. 4C, D). For example, TFAM and ssDNA formed droplets with ssDNA at the periphery of the droplet in the simulations, whereas ssDNA and mtSSB formed a droplet with ssDNA at the core of the droplet, as seen in *in vitro* experiments. Representative simulation snapshots of these binary mixtures illustrate how each component assembled within the condensate, with protein molecules in half of the droplet rendered transparent to reveal the internal DNA organization (Fig. 4C). These spatial preferences were further quantified using radial density profiles, which report how each component was distributed from the center of the condensate to its edge (Fig. 4D).

Based on the interaction parameters we established (Fig. S4G), we next simulated the quaternary mixture of the four components: TFAM, mtSSB, dsDNA, and ssDNA. Upon simulation of this quaternary mixture, we observed that all four components were recruited into a single droplet (Fig. 5C), which further corroborated the notion that physiological mt-nucleoids exist as a phase separated entity^20,31^. We quantified the local mixing within these condensates using the radial distribution function (*g(r)*) for all pairs of components. From the first peak of *g(r),* we found that the cognate pairs of interactions for TFAM-dsDNA and mtSSB-ssDNA were still preferred contacts in the quaternary mixture over the non-cognate TFAM-ssDNA and mtSSB-dsDNA pairs of interactions (Fig. 5D). Finally, we quantified the spatial localization of these components within the condensate by calculating their radial density profile from the center of the droplet. These profiles showed that ssDNA was assembled at the core of the droplet, whereas the dsDNA enveloped the ssDNA (Fig. 5E). We also evaluated the radial distribution of dsDNA and ssDNA in the condensate formed in vitro with four components system and observed that the relative localization of these nucleic acids is similar to ones observed through coarse-grained simulations (Fig. S4E). Together, these results demonstrate that mt-replication machinery self-organizes into a multiphasic condensate whose internal structure spontaneously segregates functional states of DNA.

### mtSSB partitions into replicated mt-nucleoids *in vivo*

To relate the observed multiphase organization *in vitro* and *in silico* to actively replicating nucleoids *in vivo*, we carried out immunofluorescence and EdU imaging of HeLa cells overexpressing TFAM-mClover (Fig. 6A). We visualized hundreds of mt-nucleoids, enriched in TFAM and mtDNA, in each cell, and detected EdU signal in approximately 20% of the mt-nucleoids, consistent with previous reports that only a fraction of mt-nucleoids is actively replicating at a given time^32^. However, we noted that mtSSB had distinct localization from the first three markers: mtSSB typically localized as puncta distinct from mt-nucleoids. Interestingly, the instances in which mtSSB associated with a mt-nucleoid corresponded to the actively replicating (EdU positive) mt-nucleoids.

**Fig. 6.**
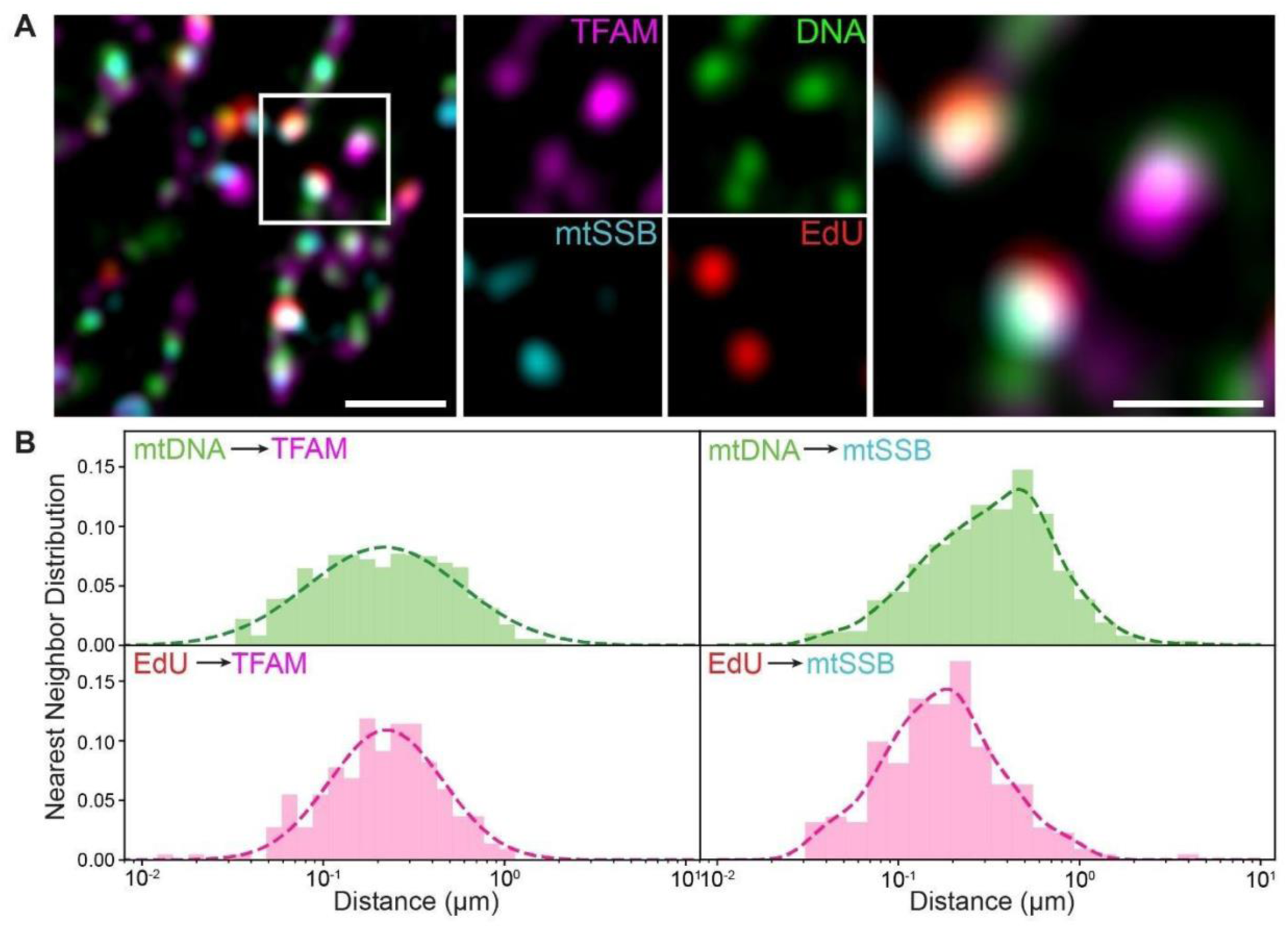
Organization of TFAM and mtDNA in active mt-nucleoids *in vivo*. A) Airyscan images of a fixed HeLa cell overexpressing TFAM-mClover (magenta), treated with EdU (red) for 1 hr before fixation, and labelled with mtDNA (anti-DNA, green) and mtSSB (anti-mtSSB, cyan). Four channel overlay with indicated white box (left, scale bar = 1 μm), individual channels (middle) and magnified version (right, scale bar = 0.5 μm). B) Probability density of nearest neighbor distance between pairs of components for mtDNA and nearest TFAM (*d_mtDNA, TFAM_*, upper left), EdU and nearest TFAM (*d_EdU, TFAM_* bottom left), mtDNA and nearest mtSSB (*d_mtDNA, mtSSB,_* upper right), and EdU and nearest mtSSB (*d_EdU, mtSSB,_* bottom right). (left: p-value = 0.88, unpaired t-test, two-tailed; right: p-value 3.2 × 10^−2^^4^, Mann-Whitney U-test, two-tailed. n = 683 for mtDNA, n= 168 for EdU).

To quantitatively measure the spatial relationship of TFAM and mtSSB puncta relative to mt-nucleoids, we performed nearest-neighbor distance analysis^33^. TFAM appeared to be closely associated with mt-nucleoids regardless of the replication state, with the average distance to the nearest TFAM punctum to all mt-nucleoids *d_mtDNA,TFAM_* = 0.30 ± 0.01 μm (mean ± s.e.m., n = 1,079 mt-nucleoids) and only replicating mt-nucleoids *d_EdU,TFAM_* = 0.26 ± 0.01 μm (mean ± s.e.m., n = 219 mt-nucleoids) (Fig. 6B). In contrast, the association of mtSSB with mt-nucleoids was strongly affected by the replication state: the average distance of the nearest mtSSB punctum to all mt-nucleoids was *d_mtDNA,mtSSB_* = 0.46 ± 0.02 μm (mean ± s.e.m., n = 1,079 mt-nucleoids), while the distance of the nearest mtSSB punctum to only replicating nucleoids (EdU positive) was significantly reduced to *d_EdU,mtSSB_* = 0.22 ± 0.01 μm (mean ± s.e.m., n = 219 mt-nucleoids), indicating high co-localization of mtSSB within 200 nm of replicating mt-nucleoids. We conclude that the localization of mtSSB to mt-nucleoids is strongly dependent on their replication activity.

### Exposed ssDNA increases mtSSB partitioning into TFAM-dsDNA rich condensates

To understand the driving force influencing mtSSB partitioning in actively replicating mt-nucleoids, we next designed single-molecule experiments with the dual optical tweezer. We began with a similar experimental approach as described above (Fig. 3), in which we tethered λ-DNA between trapped beads and exposed it to 500 nM TFAM to promote condensate formation on the DNA strand. We then moved the tethered TFAM-DNA condensate to a channel containing dilute mtSSB. We then proceeded to perform a series of stretch-relax cycles in the presence of soluble mtSSB. For the first stretch, there was no detectable mtSSB in the condensate, and we observed a similar force-extension curve as previously shown for only TFAM-DNA condensates (Fig. 7C). However, upon relaxation and additional stretch-relaxation cycles, we noticed a marked shift in the force extension curve that corresponded to increasing amounts of mtSSB partitioning within the tethered TFAM-DNA condensate (Fig. 7A, B, Supplementary Video S2). After several cycles, the force-distance relationship exhibited a significant reduction in the amount of force for a given distance and lacked a plateau in the force, reminiscent of force-extension behavior characteristic of ssDNA^34^. This observation indicated that the original dsDNA strand likely underwent force-induced melting, resulting in the *de novo* formation of ssDNA. Due to the greater flexibility of ssDNA compared to dsDNA, the DNA exhibited increased extension under the same applied force, consistent with the increasing amount of ssDNA generated with each cycle^35^. As a control, we repeated the experiment without the stretch-relax cycles and saw minimal accumulation of mtSSB within the condensate within the same amount of time (Fig. S6E, Supplementary Video S3), indicating that mtSSB accumulation was largely dependent on the presence of ssDNA. Together, these results support that only upon production of ssDNA is mtSSB able to partition within a TFAM-mtDNA co-condensate.

**Fig. 7.**
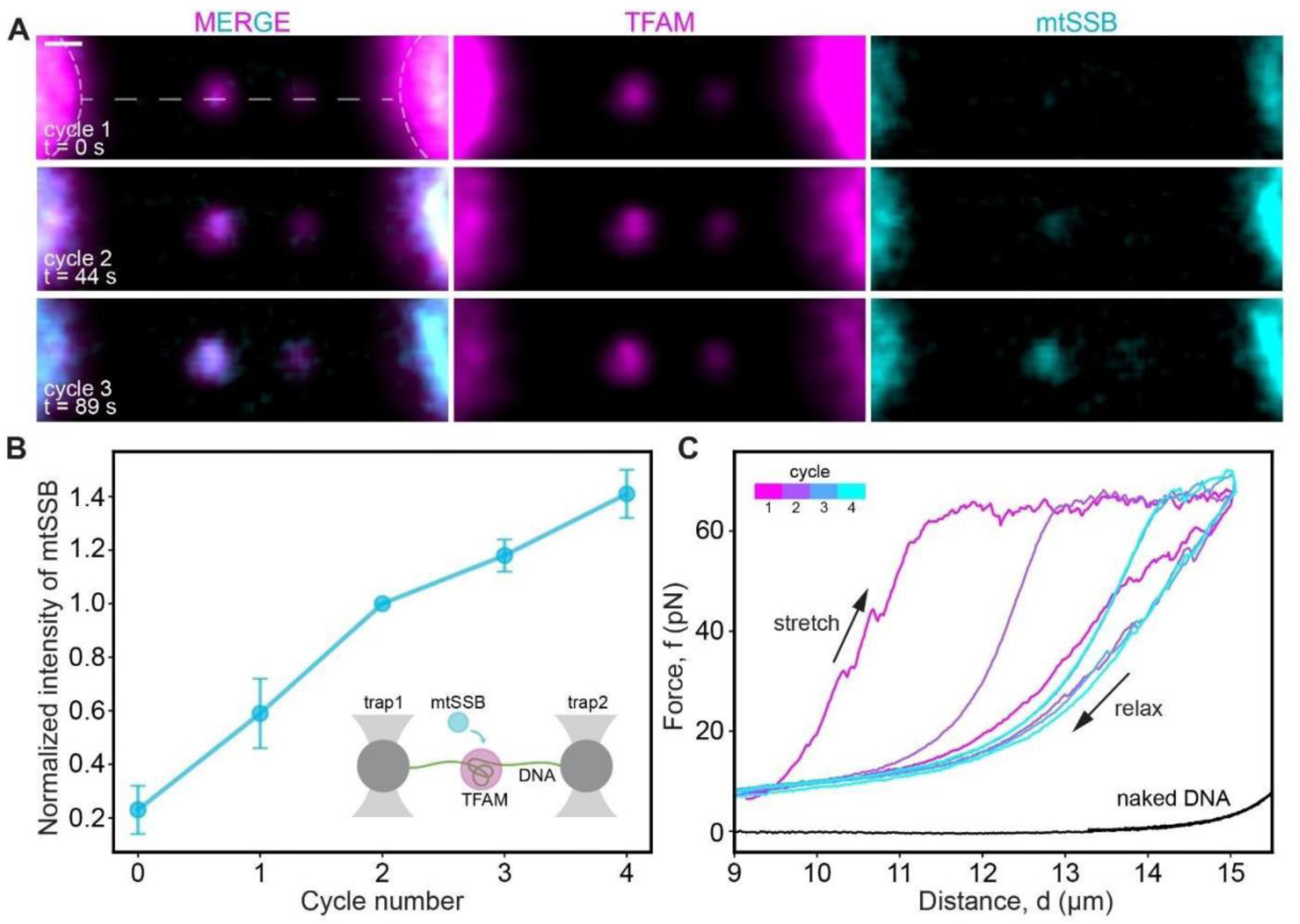
mtSSB partitioning into TFAM-DNA condensates upon repeated cycles of stretching and relaxation. A) Confocal images of a condensate (arrowhead) formed on tethered DNA (dashed line) between two optical traps (left, right) during three consecutive stretch-relaxation cycles (TFAM in magenta, mtSSB in cyan). Scale bar = 1 µm B) Normalized intensity within a condensate as a function of cycle number. Inset: schematic of the single-molecule experiment in which TFAM (magenta) first condensed on tethered λ-DNA followed by incubation with mtSSB (cyan) and several stretch-relaxation cycles. C) Force-distance curves of stretch-relax cycles of λ-DNA with TFAM condensates after incubation with mtSSB. Curves in magenta-cyan represent increasing cycle number. Black curve is naked DNA (control).

## Discussion

We show here that the core biopolymers associated with mitochondrial replication self-assemble into multiphasic condensates. We find each architectural protein spontaneously associates with its cognate nucleic acid, effectively organizing replication into sub-phases within a condensate. These results support a mechanism by which mtSSB partitions specifically into replicating mt-nucleoids, as ssDNA strands are actively generated during replication.

Our *in vitro* reconstitution allowed us to identify the relevant molecular driving forces that support the higher-order structure of replicating mt-nucleoids. We first found that differences in TFAM–DNA binding affinity, arising from DNA sequence features, such as promoter regions, led to heterogeneous internal organization within condensates. Stronger TFAM–DNA interactions, as expected for promoter regions of mtDNA, thus appear to drive local DNA compaction and promote spatial segregation of TFAM-rich and DNA-rich regions, providing a physical basis for condensate heterogeneity. Moreover, we further found length-dependent effects on organization and overall morphology: TFAM and dsDNA had scaffold-like roles on short and long length scales, respectively.

Collectively, TFAM-DNA interactions gave rise to a condensate that sustains high forces (>10 pN), strongly resisting the unpackaging of dsDNA. Such forces are considerably higher than those reported in other systems, such as for heterochromatin protein 1 (HP1a)^10^, Fox1A^36^ or FUS^27^, needed to compact DNA. Such high packaging forces are physiologically relevant as replicative DNA polymerases (DNAPs) often need to overcome mechanical barriers due to factors such as DNA torsion or supercoiling during the process of replication^37^. A wide range of DNAPs can continue replicating against stall forces of roughly 10–40 pN^38^. Among these DNAPs, the bacteriophage polymerase T7, sharing significant structural and functional similarity to the mitochondrial DNA polymerase, POLγ, is able to continue replicating against as high as ∼30 pN stall forces^39^. The high forces associated with the condensate would thus be able to withstand the forces exerted upon replication, potentially allowing condensates to help ensure robust organization of the mt-genome during replication.

When we combined the four core replication machinery components *in vitro*, we observed complete segregation of ssDNA and dsDNA within the same condensate. Through coarse-grained simulations, we showed that the interaction strengths determined from binary mixtures are sufficient to recapitulate the assembly of the quaternary mixture. We thus conclude that the complex behavior of this quaternary mixture is determined by the additive effects of its binary components.

Within the resulting heterogenous droplet, we find that the interactions between cognate pairs of the protein and DNAs remain intact, highlighting the possibility that multiphase behavior is conducive to replication. This core-shell architecture has clear biological relevance. Localization of ssDNA together with mtSSB at the condensate core likely stabilizes vulnerable single-stranded replication intermediates and provides a protected environment for ongoing DNA synthesis. In contrast, enrichment of dsDNA and TFAM toward the outer regions may facilitate genome compaction, mechanical stability, and accessibility for transcription or remodeling. Interestingly, the spatial separation of single-stranded and double-stranded nucleic acid species in condensates has been demonstrated in other contexts, particularly for single and double-stranded RNA to different phases in multiphasic peptide condensates^40^. Our results thus support a general mechanism by which the nucleic acids can be segregated based on their strandedness.

Through our fixed cell experiments, we identified that mtSSB does not associate with the majority of mt-nucleoids, but rather is excluded and forms independent puncta. However, for select mt-nucleoids, we observed significant mtSSB partitioning. Several reports identified a small fraction (∼10%) of mt-nucleoids to be actively replicating^32^. Indeed, we found a large fraction of actively replicating mt-nucleoids (70% of EdU positive nucleoids) to be enriched in mtSSB, compared to a minority inactive mt-nucleoids (36% of EdU negative nucleoids), suggesting the difference in mt-nucleoid replication activity largely determines mtSSB partitioning. Our single-molecule experiments provide direct evidence that ssDNA increases the solubility of mtSSB with TFAM-dsDNA rich condensate due to the observed partitioning of mtSSB upon force-induced melting of dsDNA. Our model becomes that actively replicating mt-nucleoids expose ssDNA, which increases the solubility of mtSSB, allowing it to accumulate within the condensate and associate with ssDNA.

These thermodynamic driving forces may be relevant for replication more broadly. For example, mtSSB is similar in structure to the *E. coli* single-stranded DNA binding protein (SSB)^41^, which has been reported to form biomolecular condensates *in vitro*^42,43^. Indeed, in *E. coli*, the majority of SSB localizes to many DNA-free membrane-proximal condensates in addition to DNA replication forks and DNA damage sites^44^. This is consistent with our observation in human mitochondria, where most mtSSB puncta were not in direct contact with mt-nucleoids. Interestingly, in the mammalian nucleus, the major nuclear ssDNA binding protein, Replication Protein A (RPA)^45^, is unrelated in structure to mtSSB or SSB, yet has also been observed to undergo phase separation, potentiated by ssDNA^46^. Our results suggested that multiphase behavior driving the segregation of ds- and ssDNA may be a fundamental organizational principle for replicating genomes. Overall, our work highlights how the thermodynamics of the core replication biopolymers determine their spatial organization, providing a physical framework for understanding how mt-nucleoids dynamically coordinate genome replication.

## Acknowledgments

We thank all the members of Feric group, Mittal group, and PSU Center for Eukaryotic Gene Regulation (CEGR) for their feedback and discussion. Research reported in this publication was supported by the National Institute of General Medical Sciences of the National Institutes of Health under award number R35 GM154931 (MF) and R35 GM153388 (JM). We thank the Genomics Core Facility (RRID:SCR_02364) for whole-genome sequencing of our plasmids. We thank the Biomolecular Interactions Core Facility (RRID:SCR_024464) for C-trap experiments (S10OD038190). We thank the Texas A&M High Performance Research Computing (HPRC) for providing computational resources required for performing the molecular dynamics simulations presented in this work.

## Resource Availability

### Lead Contact

Requests for further information and resources should be directed to and will be fulfilled by the lead contact, Marina Feric.

### Materials availability

Plasmids generated in this study are available upon request to Marina Feric.

### Data and code availability

- Original images as part of this work have been deposited to the Feric Lab GitHub repository and is publicly available via https://github.com/fericlab/mt-replication-condensates.
- All original code developed as part of this work for simulations and their analysis have been deposited to a GitHub repository and is publicly available via https://github.com/tangade-ashish/Minimalistic-Model-for-simulation-of-Protein-DNA-condensates
- Any additional information required to reanalyze the data reported in this paper is available from the lead contact upon request.

### Author contributions

M.F, J.M. and Y.Y. designed the study. Y.Y. performed *in vitro* and EdU/immunofluorescent experiments and analyzed data. Q.L. synthesized ssDNA and performed binary/quaternary mixing experiments. N.P. performed tissue culture experiments. S.T.P. wrote image analysis code. A.S.T performed simulations. A.S.T and A.M analyzed the simulation data. All authors reviewed the manuscript.

### Limitation of the study

In this study, we observed similar organization of mitochondrial replication components *in vitro*, *in vivo* and *in silico*, suggesting our model captured the main driving forces underlying the co-condensation of mt-replication components. In doing so, we designed a highly simplified *in vitro* system consisting of only of the core architectural molecules (dsDNA, ssDNA, TFAM, and mtSSB), meaning we did not actually initiate replication *in vitro*. Future studies would need to consider the contributions from active DNA synthesis and how additional replication components (POLG1/2, TWINKLE) may further influence component interaction strengths and overall condensate organization. Moreover, we used linear segments of the mt-genome for our high-resolution imaging and lambda-phage DNA for our optical tweezer experiments; we do not expect the deviations from the endogenous circular mtDNA to be significant, as TFAM and mtSSB predominantly bind DNA non-specifically and locally. For *in vivo* experiments, our study only imaged a single cell line (HeLa) that overexpressed fluorescently labelled TFAM. Future studies of colocalization of mt-replication components across different cell lines and in different functional contexts would expand our understanding structure-function relationships underlying mitochondrial replication.

### Declaration of generative AI

ChatGPT (OpenAI) and Claude (Anthropic) were used to assist with writing code for simulations and image analysis as indicated in the methods. During the preparation of this manuscript, the authors also used ChatGPT (OpenAI) and Claude (Anthropic) for improving grammar and writing clarity. All output generated by the tool were subsequently reviewed and edited by the authors, who take full responsibility for the final content of the publication.

## STAR Methods

### Key resources table

| REAGENT or RESOURCE | SOURCE | IDENTIFIER |
| --- | --- | --- |
| **Antibodies** |  |  |
| Rabbit polyclonal anti-SSBP1 | Sigma | Cat#HPA002866; RRID: AB_1080092 |
| Mouse monoclonal anti-DNA | Millipore | Cat#CBL186; RRID: AB_11213573 |
| Goat anti-rabbit IgG (H+L), Alexa Fluor 568 | Thermo Scientific | Cat# A-11036; RRID: AB_10563566 |
| Goat anti-mouse IgG (H+L), Alexa Fluor 405 | Thermo Scientific | Cat# A-31553; RRID: AB_221604 |
| **Bacterial and virus strains** |  |  |
| DH5α | New England Biolabs | C2987 |
| BL21 DE3 | New England Biolabs | C2527H |
| **Chemicals, peptides, and recombinant proteins** |  |  |
| Abberior STAR RED NHS carbonate | Abberior | Cat#STRED-0002-1MG |
| Dylight™ 488 NHS ester | Thermo Scientific | Cat#46403 |
| Poly(ethylene glycol), average M<sub>N</sub> 3,350 | Millipore | Cat#202444 |
| Biotinylated Lambda DNA | LUMICKS | Cat#00001 |
| **Critical commercial assays** |  |  |
| TaKaRa LA Taq DNA Polymerase | Takara Bio | Cat#RR002A |
| Quick-Load Taq 2X Master Mix | New England Biolabs | Cat#M0271L |
| GeneJET PCR Purification Kit | Thermo Scientific | Cat#K0701 |
| Zymoclean Large Fragment DNA Recovery | Zymo Research | Cat#D4045 |
| GeneJET Gel Extraction Kit | Thermo Scientific | Cat#K0691 |
| ULYSIS™ Alexa Fluor™ Nucleic Acid Labeling Kit | Thermo Scientific | Cat#U21654 |
| Click-iT™ Plus EdU Cell Proliferation Kit for Imaging, Alexa Fluor™ 647 dye | Thermo Scientific | Cat#C10640 |
| **Deposited data** |  |  |
| Image analysis code | This paper | https://github.com/fericlab/mt-replication-condensates |
| Coarse grained simulation code | This paper | https://github.com/tangade-ashish/Minimalistic-Model-for-simulation-of-Protein-DNA-condensates |
| **Experimental models: Cell lines** |  |  |
| HeLa | ATCC | Cat#CCL-2; Lot#70046455; RRID: CVCL_0030 |
| hTert-CRL-1474 | Feric et al.<sup>20</sup> |  |
| **Oligonucleotides** |  |  |
| Primers for PCR; See Table S1, S2 | This paper | N/A |
| Recombinant DNA |  |  |
| Human_TFAM_NoMTS_Pet28 | Addgene | RRID: Addgene_34705 |
| pET-28a+_mtSSB | This paper | N/A |
| pcDNA3 TFAM-mClover | Addgene | RRID: Addgene_129574 |
| Software and algorithms |  |  |
| ImageJ (1.54f) | Schneider et al.<sup>47</sup> | https://imagej.nih.gov/ij/ |
| HOOMD-blue (4.7.10) | Anderson et al.<sup>48</sup> | https://hoomd-blue.readthedocs.io/en/stable/index.html |
| Visual Molecular Dynamic (VMD, 2.0.0a4) | Humphrey et al.<sup>49</sup> | https://www.ks.uiuc.edu/Research/vmd/<br>RRID:SCR_001820 |
| Freud library | Ramasubramani et al.<sup>50</sup> | https://freud.readthedocs.io/en/stable/index.html |
| Zen Blue (3.6.095.03000) | Carl Zeiss | RRID: SCR_013672 |
| Zen Black (3.0 SR FP2) | Carl Zeiss | RRID: SCR_013672 |
| Adobe Illustrator 2025 | Adobe | https://www.adobe.com/products/illustrator.html<br>RRID:SCR_010279 |
| Other |  |  |
| HiTrap Heparin HP affinity column | Cytiva | Cat#17040701 |
| Amicon Ultra Centrifugal Filter, 10 kDa MWCO | Millipore | Cat#UFC501008 |
| Amicon Ultra Centrifugal Filter, 3 kDa MWCO | Millipore | Cat#UFC500308 |
| Grace Bio-Labs Press-To-Seal silicone isolator, PSA both sides | Millipore | Cat#GBL666105 |
| TetraSpeck™ Microspheres, 0.1 µm, fluorescent blue/green/orange/dark red | Thermo Scientific | Cat#T7279 |
| Streptavidin-coated 4.0-4.9 µm beads | LUMICKS | Cat#11228 |

