## Supplementary Information for "Organization of core mitochondrial replication components into multiphasic condensates"

Supplemental Information

**This PDF file includes:**

Methods

Tables S1 and S2

Supplemental References

Extended Data Figures S1 – S6

Supplementary Video Legends

**Other Supplementary Materials for this manuscript include:**

Supplementary Videos 1 to 3

### Methods

#### Protein purification

Mature human TFAM (43-246) Addgene plasmid #34705) and mtSSB (17-148) (GenScript, MHS6278-202756022) proteins were purified using pET28a+ expression vectors. Briefly, BL21 DE3 pRare *E. coli* was transformed with each construct. A bacterial culture (5L) of Dynamite media plus kanamycin was inoculated and incubated until an OD600 of 0.7 was reached. IPTG was added at 0.5 mM to induce protein expression, and the culture was incubated overnight at 16°C at 220 rpm. The culture was harvested with a Sharples T-1-P Super Centrifuge and the bacterial pellet was resuspended and lysed in Lysis Buffer (20 mM Tris-HCl, 500 mM NaCl, 25 mM imidazole, pH 8.0). Cells were mechanically lysed with a LM-20 microfluidizer at 25,000 psi and centrifuged at 38,000 g for 30 minutes at 4°C. Proteins were purified using a fast protein liquid chromatography (FPLC) immobilized metal affinity chromatography (IMAC) column. For TFAM, the protein was eluted with Elution Buffer 1 (20 mM Tris-HCl, 500 mM NaCl, 100 mM imidazole) then Elution Buffer 2 (20 mM Tris-HCl, 500 mM NaCl, 250 mM imidazole). Fractions containing TFAM were pooled and dialyzed into Storage Buffer (20 mM Tris, 500 mM NaCl, pH 8.0) and stored at 4°C. For mtSSB, the protein was eluted with Elution Buffer 2 and filtered into lysis buffer through tangential flow filtration.

To remove any bound nucleic acids, a 5 mL HiTrap Heparin HP affinity column (Cytiva) was used. For TFAM, the protein was first diluted in Heparin Buffer 1 (20 mM Tris-HCl, 300 mM NaCl, pH 7.5). The column was pre-equilibrated with Heparin Buffer 1; the protein was loaded onto the column; the column was washed with 10X column volumes of Heparin Buffer 1. The protein was eluted with a two-step gradient of 700 mM NaCl followed by 1 M NaCl. Fractions containing purified protein were pooled together. For mtSSB, the protein was diluted in Heparin Buffer 2 (20 mM Tris-HCl, 50 mM NaCl, pH 7.5). The column was pre-equilibrated with Heparin Buffer 2; the protein was loaded onto the column; the column was washed with 10X column volumes of Heparin Buffer 2. The protein was eluted with a two-step gradient of 700 mM NaCl followed by 1 M NaCl. Fractions containing purified protein were pooled together, resuspended in 10% (v/v) glycerol, frozen in liquid nitrogen and stored at -80°C.

#### Protein labeling

To fluorescently label TFAM and mtSSB, ~0.3 mg of each protein was dialyzed into PBS. 100 uL of 0.2 M sodium bicarbonate was added per 1 mL of protein solution (0.1 – 0.3 mg/ml). STAR-RED NHS carbonate (Abberior) or Dylight 488 (Thermo Scientific) was dissolved in DMSO to a concentration of 10 mg/mL. Fluorescent dye was added to the protein solution to a final concentration of ~0.5 mg/ml for one hour at room temperature under constant rotation and protected from light. After the reaction, the labeled protein was dialyzed into 20 mM Tris, 500 mM NaCl, pH 7.5. Glycerol was added to a final concentration of 10% (v/v). Labelled protein was subsequently frozen in liquid nitrogen and stored at -80°C.

To determine the labelling efficiency of proteins, absorption at 493 nm (for mtSSB-Dylight 488) or 638 nm (for TFAM-STAR RED) and at 280 nm was measured using a Thermo Fisher Nanodrop

One. Using known extinction coefficients for each dye, the labeling efficiency was calculated using the equation below:

$$\frac{Dye}{protein} = \frac{A_{dye} \times \epsilon_{protein}}{(A_{280} - A_{dye} \times CF_{280}) \times \varepsilon_{dye}}$$

We obtained labelling efficiencies of ~0.9 molecules of dye per protein and ~0.8 molecules of dye per protein for TFAM-STAR RED and mtSSB-Dylight 488, respectively.

**dsDNA amplification and purification**

To obtain mitochondrial DNA, two overlapping fragments of the mtDNA were amplified from genomic DNA by long-range PCR with Takara LA Taq DNA polymerase using two sets of primers (~9 kb fragment: 5'-AAC CAA ACC CCA AAG ACA CC-3' and 5'-GCC AAT AAT GAC GTG AAG TCC -3' and ~7 kb fragment: 5'-TCC CAC TCC TAA ACA CAT CC-3' and 5'-TTT ATG GGG TGA TGT GAG CC-3')[1].

To test the contribution of sequence features, three 100 bp DNA fragments were amplified by PCR (see Table 1) using the mtDNA fragments as templates. Sequences containing the light strand promoter (LSP) and heavy strand promoter (HSP) were amplified from the 7 kb fragment, while a non-specific (NS) sequence was amplified from the 9 kb fragment. To test the contribution of DNA length, DNA fragments of 0.2, 0.5, 1, 2, and 5 kb were amplified by PCR using the 9 kb mtDNA fragment as the template (see Table 2). The 9 kb template was used to approximate 10 kb. All DNA templates were purified with PCR clean-up kits and eluted in nuclease-free water.

**dsDNA fluorescent labelling**

The 100 bp DNA fragments were fluorescently labeled with Alexa Fluor 594 using Ulysis nucleic acid labeling kit (Invitrogen). 1 ug of amplified DNA was diluted with 20 uL of labeling buffer. 5 uL of ULS labeling reagent stock solution was added to the DNA and allowed to react at 80°C for 15 minutes. The temperature then was slowly dropped to 25°C. The labeled DNA was recovered using a P30 column or PCR purification kit, as purchased. The labeling efficiency of DNA was measured by measuring absorption at 260 nm and 590 nm. Based on known extinction coefficients, labeling efficiency was calculated using equation below:

$$\frac{base}{dye} = \frac{(A_{260} - A_{590} \times CF_{260}) \times \varepsilon_{dye}}{A_{590} \times \varepsilon_{base}}$$

We obtained labelling efficiencies of ~3, 0.5, and 2 molecules of dye/molecule for HSP, LSP, NS dsDNA, respectively. The labeled DNA fragments were mixed with unlabeled counterparts to the same fluorophore to achieve the same ratio of unlabeled: labeled DNA of 0.1 dye/strand. To visualize the DNA of different lengths, DNA was stained with DAPI (1:10,000, Thermo Scientific) in solution.

**ssDNA synthesis, purification, and fluorescent labelling**

The full *Cytb* gene from the human mitochondrial genome (1,141 bp) was first amplified by traditional PCR using Quick-Load *Taq* 2X Master Mix using the 7 kb mtDNA fragment as a template with primers (Forward: 5'-ATGACCCCAATACGCAAAAC-3', Reverse: 5'-AGGCCCATTTGAGTATTTTGTT-3'). Reaction was performed at 95ºC for 30 s, 30 cycles of (i) 95ºC for 30 s, (ii) 49ºC for 30 s, (iii) 68ºC for 70 s, followed by 68ºC for 5 mins and chilled at 4ºC. The dsDNA *Cytb* product was then purified using ThermoScientific GeneJET PCR Purification Kit. Concentration was measured using a ThermoScientific Nanodrop One.

To synthesize ssDNA, we used reiterative PCR (rPCR) using only the forward primer[2]. The rPCR reaction consisted of: the dsDNA *Cytb* PCR product, only forward primer (5'-ATGACCCCAATACGCAAAAC-3'), and Quick-Load *Taq* 2X Master Mix. To generate fluorescently labelled ssDNA product, we performed the rPCR reactions with a directly labelled primer (5'- /5RhoR-XN/ ATGACCCCAATACGCAAAAC-3', IDT), ensuring that each ssDNA transcript had a single fluorophore at the 5' end. Reactions were performed at 95ºC for 30 s, 30 cycles of (i) 95ºC for 30 s, (ii) 49ºC for 30 s, (iii) 68ºC for 70 s, followed by 68ºC for 5 mins and chilled at 4ºC. Total product was first purified and concentrated using ThermoScientific GeneJET PCR Purification Kit. Next, the resulting concentrated product containing original template dsDNA and synthesized ssDNA product was run on a 1% agarose gel stained with SYBR Safe followed by gel extraction. To isolate the ssDNA, the lower molecular weight ssDNA band was specifically excised and purified using ThermoScientific GeneJET Gel Extraction Kit.

**TFAM and dsDNA co-phase separation assays *in vitro***

Three buffers were prepared and filter-sterilized prior to the experiments: high salt buffer (20 mM Tris–HCl, 500 mM NaCl, 10 mM $MgCl_2$, pH 7.8), low salt buffer (20 mM Tris–HCl, 10 mM $MgCl_2$, pH 7.8), and low salt buffer with PEG (20 mM Tris–HCl, 10 mM $MgCl_2$, 30% PEG-3K (Sigma-Aldrich), pH 7.8). Frozen aliquots of protein were thawed at room temperature and then stored on ice. Protein solutions were concentrated and buffer exchanged (20 mM Tris–HCl, 500 mM NaCl, 10 mM $MgCl_2$, pH 7.8) using 0.5 ml centrifugal filters 10-kDa (Amicon) and a tabletop centrifuge at 4°C. Final protein concentration was measured by Bradford Assay using BSA standards (Bio-Rad) on a spectrophotometer (GENESYS 50 UV-Vis Spectrophotometer, Fisher Scientific). Protein solution was mixed with varying concentration of buffers to reach the indicated concentrations. Specifically, buffers were mixed first, followed by mtDNA and lastly protein. Solutions were then mixed by gently flicking and briefly centrifuged prior to imaging. The final ratio of TFAM to DNA was 1 TFAM molecule : 6 bp DNA.

**Four component phase separation assays *in vitro***

Frozen protein aliquots of mtSSB and TFAM were thawed at room temperature. Protein solutions were then concentrated and buffer exchanged (20 mM Tris-HCl, 500 mM NaCl, 10 mM $MgCl_2$, pH 7.8) using 3K or 10K Amicon filters, respectively, and a tabletop centrifuge at 4ºC. Final protein concentrations were measured by Bradford Assay using BioRad BSA Standards on a ThermoScientific Genesys 50 spectrophotometer. Fluorescently labeled units of mtSSB, TFAM, and ssDNA were combined with unlabeled units in a 1:10 ratio of labeled : unlabeled. dsDNA was visualized with DAPI (1:10,000). Concentrated protein solutions were combined with low salt buffer (20 mM Tris-HCl, 0 mM NaCl, 10 mM $MgCl_2$, pH 7.8) and low salt buffer containing PEG to yield a final solution containing 20 µM TFAM, 20 µM mtSSB, 20 nM ssDNA and/or 20 nM dsDNA in 20 mM Tris-HCl, 150 mM NaCl, 10 mM $MgCl_2$, 5% PEG, pH 7.8. Order of addition was first buffer solutions, followed by nucleic acids, and lastly protein solutions to make a 5 µl final solution. Each mixture was spun on a tube rotator at room temperature for 30 mins. After gentle rotation, samples were transferred to an imaging chamber enclosed by a silicone isolator (Grace-Biolabs) and incubated at room temperature for an additional 30 mins prior to imaging.

**Cell Culture**

HeLa cells (ATCC, CCL-2, Lot #70046455) were cultured in Dulbecco's Modified Eagle Medium (DMEM, Thermo Fisher Scientific, #11960044) supplemented with 10% fetal bovine serum (FBS, Thermo Fisher Scientific, #A5256701) and 1% penicillin-streptomycin-glutamine (Thermo Fisher Scientific, #10378016) at 37°C with 5% $CO_2$.

**Transient transfection and EdU incorporation**

Cells were transfected with TFAM-mClover (a gift from Stephen Tait, Addgene #129574)[3] ~24 hours after seeding on coverslips in 12-well plates using FuGene HD Transfection Reagent (Promega, #E2311) following the manufacturer's optimized instructions. After ~48 hours of transfection, cells were rinsed with PBS and incubated with 20 µM EdU (from Thermo Fisher Scientific, #C10640) for 30 mins in complete media containing DMEM without phenol red (Thermo Fisher Scientific #31053028). Following EdU incubation, cells were rinsed with PBS and labelled with 100 nM MitoTracker Red in cell culture media for 30 min, fixed with 4% PFA for 20 minutes, and processed for EdU labeling and immunofluorescence (see below).

**EdU labelling and immunofluorescence**

Fixed cells were permeabilized with 0.5% Triton X-100 in PBS for 10 minutes at room temperature (RT). EdU labeling with Alexa Fluor 647 was performed using Click-iT Plus EdU Labeling Kit (Invitrogen) following manufacturer's protocol. After the labeling, cells were washed twice with PBS and a second permeabilization was performed with 0.5% Triton X-100 in PBS for 10 minutes at room temperature. Cells were then washed and incubated with primary antibodies in PBS with 5% BSA for 1 hour at RT: anti-SSBP1 (Sigma, HPA002866, 1:1,000 dilution) and anti-DNA (Millipore, CBL186, 1:500 dilution). Cells were washed three times with PBS followed by incubation with secondary antibodies for 1 hour at room temperature: anti-mouse-Alexa Fluor 405 (Invitrogen, A31554, 1:50 dilution), and anti-rabbit-Alexa Fluor 568 (Invitrogen, A11036, 1:1,000 dilution). Cells were washed three times with PBS and incubated in 4% PFA in PBS for 10 minutes at room temperature. Coverslips were mounted onto glass slides with ProLong Gold Antifade Mountant (Thermo Scientific) and left to cure for 48 h.

### Light microscopy

High-resolution imaging was performed using a laser scanning confocal microscope equipped with an Airyscan 2.0 detector (Zeiss LSM 980 series). A Plan-Apochromat 63x/1.40 oil DIC M27 oil immersion objective was used to collect images. Raw images were processed through the Airyscan processing function in Zen Blue software to achieve super-resolution images. Channels were aligned by calibrating with four-color beads (Thermofisher, #T7279). Slides containing 100 nm four-color beads were imaged with same setting (zoom factor, laser power, etc) used for data collection. The beads images were then subjected to fitted channel alignment in Zen Black. The alignment parameters were then saved and applied to all collected images. Confocal imaging was performed using the Abberior STEDYCON add-on module. An alpha Plan-Apochromat 100X/1.46 Oil Iris M27 objective was used for collecting images.

### Optical tweezer

Single-molecule experiments were performed at room temperature on a LUMICKS C-trap with a custom five-channel flow cell. First, streptavidin-coated polystyrene beads (4.35 μm, LUMICKS) were applied to Channel 1 and captured by the optical traps at a stiffness of ~0.4 pN/nm. Next, the traps were moved to Channel 2, which contained biotinylated λ-DNA (LUMICKS). The DNA was subsequently tethered to the two beads. The optical traps were then moved to Channel 3 for force calibration of naked DNA.

To perform force-distance measurements in the presence of TFAM condensates, biotinylated λ-DNA was first moved to Channel 4 containing 500 nM TFAM (STAR RED) in 20 mM Tris, 10 mM $MgCl_2$, 150 mM NaCl, pH 7.8 to allow TFAM condensates to form on λ-DNA. Once a condensate formed on the tethered DNA, the flow was stopped, and a force-distance measurement was carried out by repeatedly stretching and relaxing the TFAM-bound DNA.

Only for co-localization experiments of DNA within TFAM condensates, 1% PicoGreen was added to Channel 4. Trapping of beads and fishing of DNA were carried out as described above. After DNA was moved to Channel 4 and TFAM condensates formed, dual color confocal imaging (488 nm and 638 nm) was carried out while labelled λ-DNA was stretched and relaxed for multiple cycles.

For force-distance experiments in the presence of mtSSB, the experimental set up was performed as above. Briefly, tethered DNA was moved to Channel 4 as to form a TFAM bound condensate. Next, the TFAM-bound DNA was then moved to channel 5, which contained 500 nM mtSSB (DyLight 594) in buffer containing 20 mM Tris, 10 mM $MgCl_2$, 150 mM NaCl, pH 7.8. Dual color confocal imaging (561 nm and 638 nm) and force measurement were carried out while λ-DNA was stretched and relaxed for multiple cycles in the presence of mtSSB.

### Quantitative image analysis

Images were visualized using ImageJ[4] (version 1.54f) and quantitatively analyzed using python (version 3.12). To characterize droplet morphology, droplets were segmented from background by applying an overall threshold based on fluorescence intensity. Objects within a defined size range were selected as droplets. Droplet area was determined by counting the number of pixels contained in a segmented droplet, and the resulting area was used to estimate the radius ($A = \pi r^2$).

To estimate degree of miscibility between pairs of components, Pearson correlation coefficient (ρ) was calculated for pairs of channels for all pixels within the droplet and averaged across all droplets per condition.

To estimate the degree of irregularity in droplet morphology, negative curvature (M.N.C) was obtained by calculating local curvatures along the contour of each droplet, followed by taking the absolute value of the mean of negative local curvature only[5].

Nearest neighbor calculation was performed as previously described[6]. The analysis was adapted for comparing four channels.

To analyze the radial distribution of ssDNA and dsDNA intensity within representative quaternary droplets, the centroid of a droplet was determined. Then a coordinate grid covering the entire droplet was generated and the pixel intensities and distance to the center were calculated, yielding a radial intensity profile. Finally, the intensity was normalized relative to the maximum value for each channel. The code was generated using AI tool (ChatGPT) and refined manually.

**Minimalistic Coarse-Grained Model**

We used our previously developed minimalistic coarse-grained model[7] in which protein molecules were modeled as a spherical bead, and the DNA was modeled as a polymeric chain. The size of a TFAM bead ($\sigma_{TFAM}$) and mtSSB bead ($\sigma_{mtSSB}$) was 50 Å, which roughly corresponds to twice the radius of gyration of both the proteins[8]. The size of the DNA ($\sigma_{DNA}$) monomer was 5 Å, which was connected by an equilibrium bond length ($r_o$) of 5.5 Å with a harmonic bond potential ($U_{bond}$) with spring constant ($K_{bond}$) of 20 $kcal/mol/Å^2$.

$$U_{bond} = K_{bond}(r - r_o)^2$$

To model the rigidity difference between ssDNA and dsDNA, a harmonic angle potential $\left(U_{angle}\right)$ is added between three consecutive monomers with spring constants $\left(K_{angle}\right)$ of 0.5 and 20 $kcal/mol/rad^2$ respectively with the equilibrium angle ($\theta_o$) as 180°.

$$U_{angle} = K_{angle}(\theta - \theta_o)^2$$

The effective non-bonded interactions between the protein, protein-nucleic acid, and nucleic acid beads were modeled with the Ashbaugh-Hatch potential[9,10]

$$U_{AH}(r) = \begin{cases} U_{LJ}(r) + \left(1 - \lambda_{ij}\right)\varepsilon\ , & r \leq 2^{1/6}\sigma \\ \lambda_{ij} U_{LJ}(r)\ , & otherwise \end{cases}$$

where, $U_{LJ}(r)$ is the Lennard-Jones potential,

$$U_{LJ}(r) = 4\varepsilon\left[\left(\frac{\sigma}{r}\right)^{12} - \left(\frac{\sigma}{r}\right)^{6}\right]$$

where, $\varepsilon$ was fixed to a value of 0.2 $kcal/mol$ and $\lambda_{ij}$ is the average hydropathy between the two interacting species. The strength of effective interactions between components was varied by changing the value of $\lambda_{ij}$. The effective interactions between the monomers of the nucleic acids were modeled as repulsive by setting $\lambda_{DNA-DNA} = -1$, except for the bonded DNA beads. The effective interactions were modeled with $\lambda_{TFAM-TFAM} = 3.5$. The interaction parameters $\lambda_{ij}$ for modeling the binary component interactions are represented in Fig. S4G, the same parameters are used throughout this work unless specifically mentioned in the text.

**Simulation Details**

Similar to the experimental ratio of one protein molecule per six DNA base pairs, we simulated protein and DNA mixtures with a ratio of one protein molecule to six DNA monomers. For the case of 100-mer dsDNA simulations, we simulated 1,000 TFAM molecules and 60 chains of 100-mer dsDNA in a cubic box with an edge length of 1,170 Å. To accommodate the longer DNA chains, we simulated 500-mer, 750-mer, 1000-mer, 1500-mer, and 2000-mer dsDNA in a cubic box with an edge length of 2,000 Å with 5,000 TFAM molecules and 30,000 dsDNA monomers. We replicated the binary experimental data with simulations with 5,000 protein molecules and 30 chains of 1,000-mer DNA in a cubic simulation box with an edge length of 2,000 Å. For the quaternary mixture simulations, we simulated 5,000 TFAM molecules, 5,000 mtSSB molecules, 30 chains of 1,000-mer dsDNA, and 30 chains of 1,000-mer ssDNA in a cubic simulation box of length 2000 Å. We performed Langevin dynamics simulations at a fixed temperature of 300 K, with the friction coefficient $\gamma_i = \frac{m_i}{t_{damp}}$; where $m_i$ is the mass of $i^{th}$component and $t_{damp}$ is set to 1,000 ps. All the simulations were performed using the HOOMD-blue molecular dynamics engine (version 4.7.0)[11] along with features from azplugins (version 0.11.0). Each simulation was run for 2 μs using a 10 fs timestep. The initial 400 ns was discarded as the equilibration period (Fig. S1C), and the remaining 1.6 μs was used for analysis, which was performed using block averaging with four components.

**Visualization of simulations**

Snapshots from the simulation trajectory are generated using Visual Molecular Dynamics 2.0 (VMD version 2.0.0a4)[12].

**Calculation of radial distribution function, *g(r)***

Using the RDF function in the density class from Freud[13] Python library (version 2.12.1), we calculated the radial distribution function, *g(r)*.

**Calculation of the radial density profiles for components**

The largest molecular cluster in the simulation box was identified using the Freud[13] Python library (version 2.12.1). A distance criterion of $1.5\sigma_{Protein}$ was applied to define molecules belonging to the same cluster. The center of mass (COM) of the largest cluster was calculated and was regarded as the origin coordinate. The distance of all simulation beads from this COM was then calculated. The distribution of these distances was binned with a width equal to the size of the beads, yielding the radial distribution of components from the cluster center. The calculation protocol for the radius of gyration of DNA chains and the protein bound to DNA is mentioned in our previous work[7].

**Calculation of the relative shape anisotropy, $\kappa^2$**

Relative shape anisotropy, $\kappa^2$, quantifies the shape of a condensate where $\kappa^2 = 0$ indicates a spherical and $\kappa^2 = 1$ indicates a cylindrical shape. We calculated the relative shape anisotropy based on the gyration tensors for the condensate as[14]:

$$\kappa^2 = 1 - 3\frac{\lambda_1\lambda_2 + \lambda_2\lambda_3 + \lambda_1\lambda_3}{(\lambda_1 + \lambda_2 + \lambda_3)}$$

where, $\lambda_i$ are the three eigenvalues of the gyration tensor.

The gyration tensor for the condensate was computed using ClusterProperties function in the Cluster class from Freud[13] Python library (version 2.12.1).

**Supplementary Table S1: primers used to generate dsDNA of different sequence features**

| | |
|---|---|
| LSP-FWD | 5'-TAA CCA GAT TTC AAA TTT TAT CTT TT-3' |
| LSP-REV | 5'-AGA TTA GTA GTA TGG GAG TGG GA-3' |
| HSP-FWD | 5'-ATA CTA CTA ATC TCA TCA ATA CAA CCC CCG-3' |
| HSP-REV | 5'-GGT GTC TTT GGG GTT TGG TTG GT-3' |
| NS-FWD | 5'-ATC CCC ATA CTA GTT ATT ATC GAA ACC A-3' |
| NS-REV | 5'-TGA GTA GGT GGC CTG CAG TAA-3' |

**Supplementary Table S2: primers used to generate dsDNA of different lengths**

| | |
|---|---|
| 200bp-FWD | 5'-TTT CGG TCA CCC TGA AGT TTA T-3' |
| 200bp-REV | 5'-GCT CGT GTG TCT ACG TCT ATT C-3' |
| 500bp-FWD | 5'-CAC AGC TCT AAG CCT CCT TAT T-3' |
| 500bp-REV | 5'-GAG AAG TAG GAC TGC TGT GAT TAG-3' |
| 1kb-FWD | 5'-CCT TCA AAG CCC TCA GTA AGT-3' |
| 1kb-REV | 5'-GGT TGC GGT CTG TTA GTA GTA TAG-3' |
| 2kb-FWD | 5'-CTC ACC ATC GCT CTT CTA CTA TG-3' |
| 2kb-REV | 5'-TGA AGG CTC TTG GTC TGT ATT T-3' |
| 5kb-FWD | 5'-TTG ACC GCT CTG AGC TAA AC-3' |
| 5kb-REV | 5'-TCC GAA GCC TGG TAG GAT AA-3' |
| 9kb-FWD | 5'-AAC CAA ACC CCA AAG ACA CC-3' |
| 9kb-REV | 5'-GCC AAT AAT GAC GTG AAG TCC -3' |

### Supplementary Figures

### Supplementary Figure 1

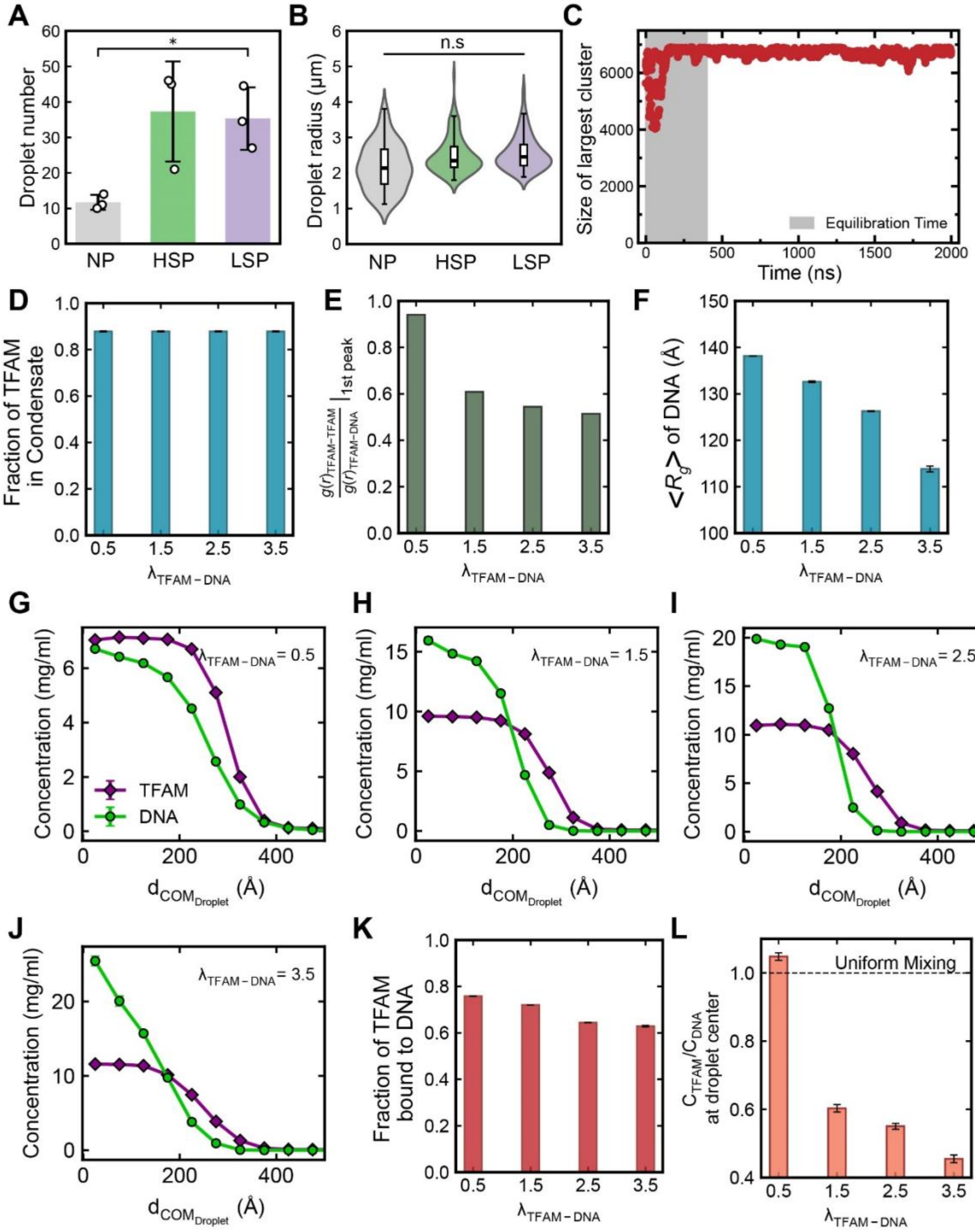

**Fig. S1 Purification and modeling of dsDNA and TFAM**

A) Droplet number in field of view. *$p < 0.05$, t-test. Error bar = S.D. n = 3 independent experimental replicates (each containing ~100s of droplets).

B) Droplet size measured from confocal images 90 min after mixing. $p > 0.05$, t-test. Error bar = S.D. n = 3 independent experimental replicates (each containing ~100s of droplets).

C) Size of the largest cluster in simulations as a function of simulation time. DNA length of 100-mer and heterotypic interaction strength of $\lambda_{PD} = 0.5$ was considered for this analysis. The initial 400 ns of the simulation was considered as an equilibration period.

D) Fraction of TFAM molecules in the condensate remains unchanged with increasing TFAM-DNA interaction strength.

E) Radial distribution function, g(r), for the TFAM & DNA where the preference of the interactions is represented by the higher peak intensity value.

F) Radius of gyration, Rg, of DNA chains in condensate decreases with increasing TFAM-DNA interaction strength indicated larger DNA collapse.

G-J) Radial density profile of the concentration of TFAM and DNA from the center of droplet for the case of 100-mer DNA at different interaction strength.

K) Fraction of TFAM bound on DNA chains in condensate decreases with an increasing interaction strength of TFAM-DNA.

L) Ratio of concentration of TFAM to concentration of DNA at the center of condensate. This ratio being 1 indicates uniform mixing of TFAM and DNA within the condensate whereas deviation from 1 indicates extent of de-mixing.

### Supplementary Figure 2

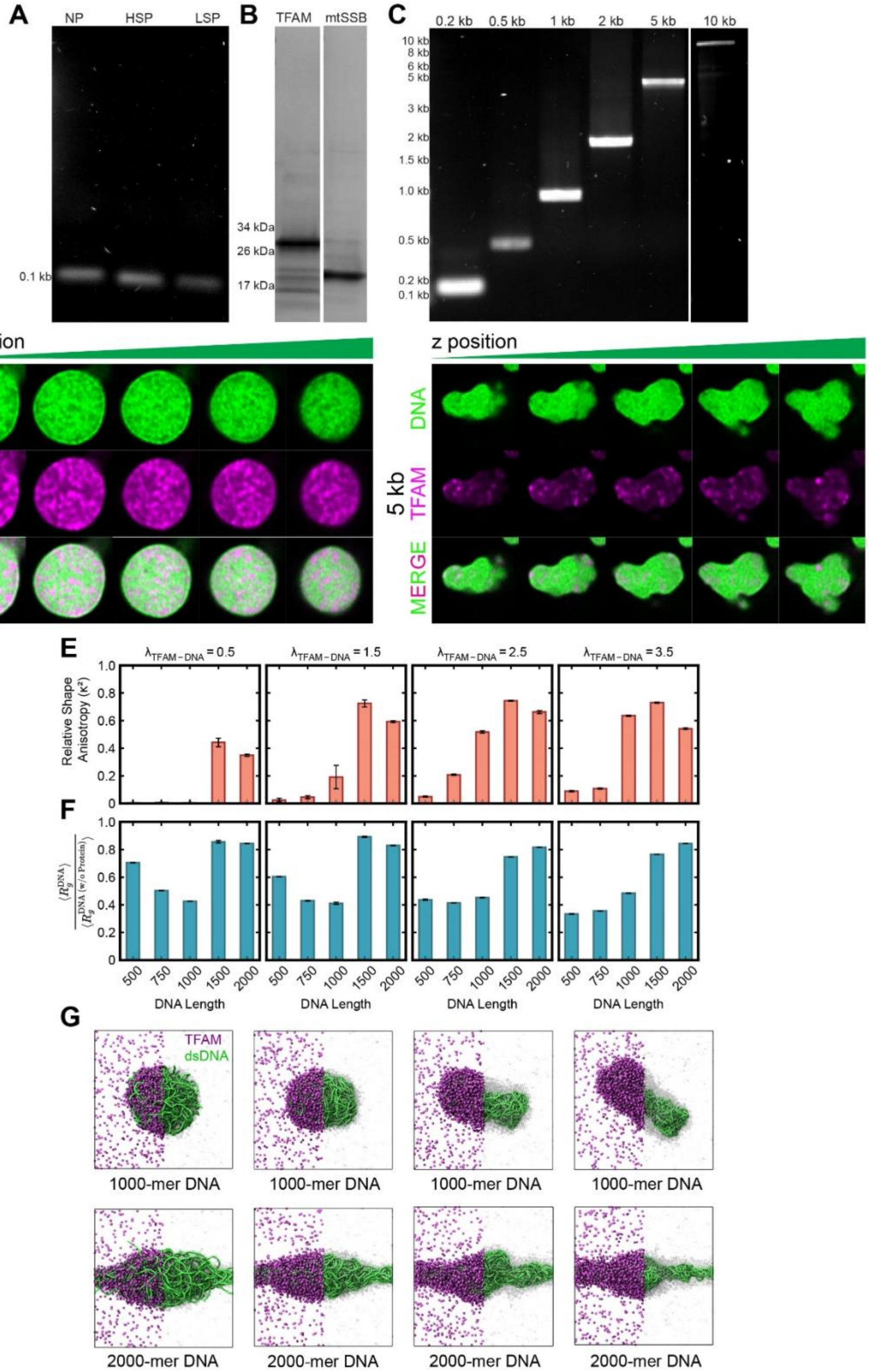

**Fig. S2 Purification and modeling of dsDNA templates of various lengths**

A) Agarose gel electrophoresis image of 100 bp DNA fragments: non-promoter (NP), heavy strand promoter (HSP), and light strand promoter (LSP).

B) Protein gel of purified 6His-TFAM and 6His-mtSSB.

C) Agarose gel electrophoresis image of non-promoter containing mtDNA fragments of varying length: 0.2, 0.5, 1, 2 and 5 and 10 kb.

D) Airyscan images of representative TFAM-mtDNA droplet with 0.5 kb and 5 kb DNA at different z positions (650 nm distance between snapshots). Scale bar = 1 μm.

E) Relative shape anisotropy for the condensate for the different interaction strengths of TFAM-DNA indicates that longer DNA forms non-spherical condensates albeit critical length required for formation of these non-spherical condensates changes with TFAM-DNA interaction strength.

F) Relative extent of DNA compaction in condensate with bare DNA without proteins computed through the minimalistic coarse-grained model.

G) Representative snapshots from the simulations highlighting the different shape of the condensate and the compaction of DNA for longer lengths of DNA.

**Supplementary Figure 3**

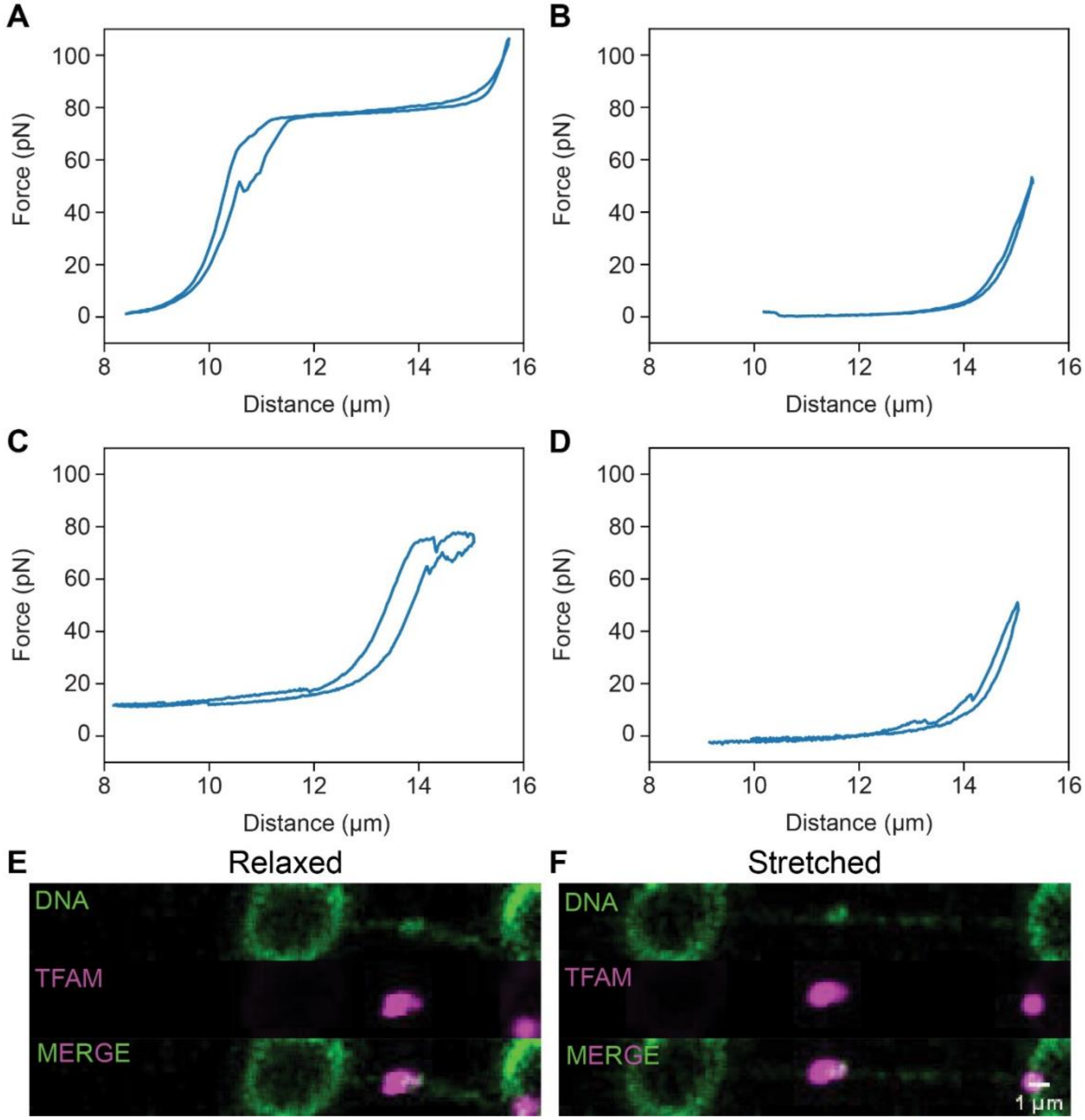


**Fig. S3 Force extension curves of TFAM-bound DNA**

A-D) Four representative force extension curves of TFAM-bound DNA using the C-trap (Lumicks). λ-phage DNA was first trapped between two beads and TFAM was applied to the flow cell channel. Stretch-relax cycle was performed on the DNA after TFAM droplet formation.

E-F) Confocal images of TFAM (magenta)-bound DNA (green, stained by PicoGreen) at (E) relaxed and (F) stretched state during pulling cycles. Scale bar = 1 μm.

**Supplementary Figure 4**

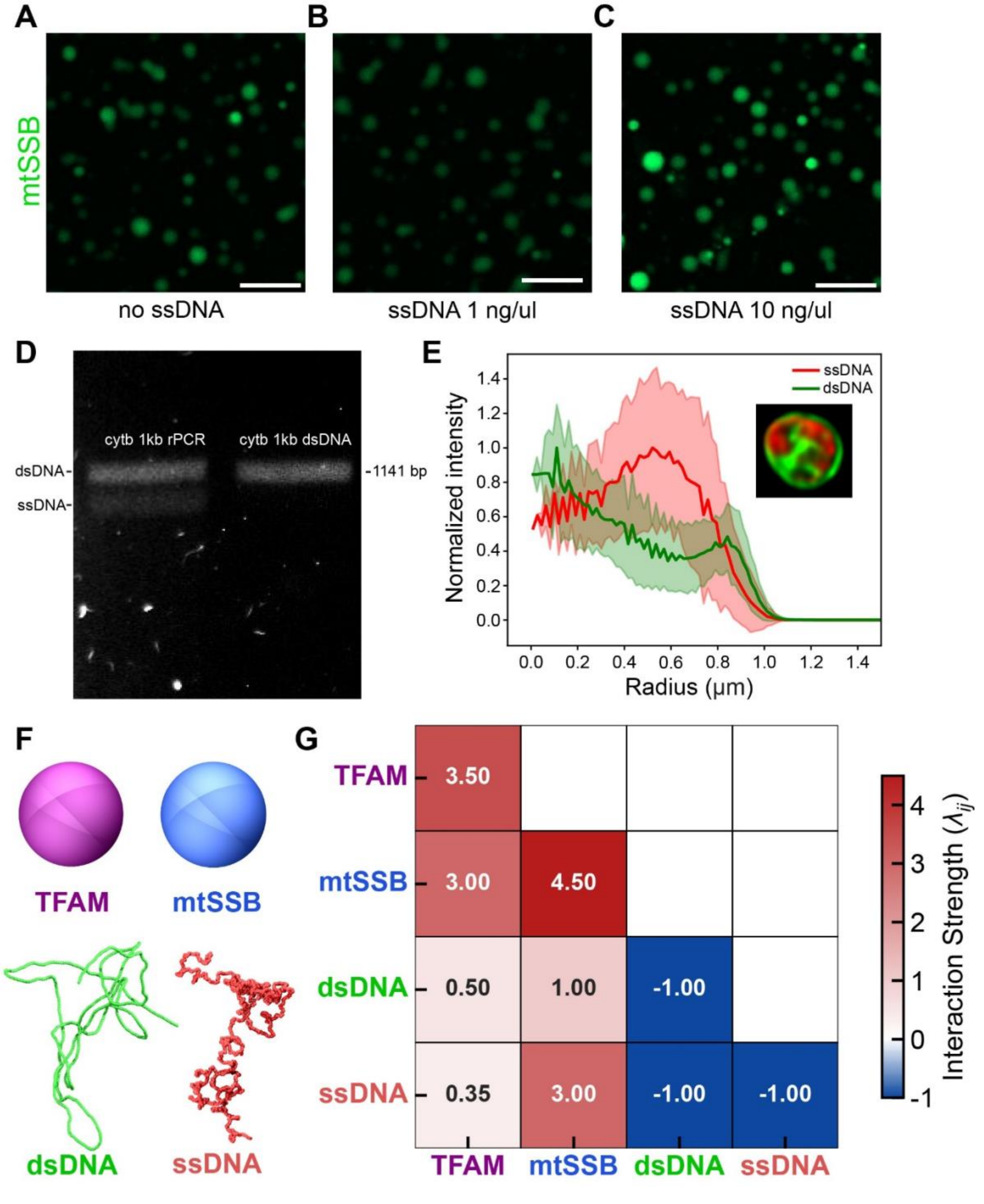

**Fig S4. Core mt-replication components spontaneously self-organize into multiphasic condensates *in vitro***

A-C) mtSSB phase separates in the absence and presence of ssDNA *in vitro*: 35 µM mtSSB (A); 35 µM mtSSB, 1 ng/ul *Cytb* 1kb sense ssDNA (B); and 35 µM mtSSB, 10 ng/ul *Cytb* 1kb sense ssDNA (C). The buffer condition for all panels is 20 mM Tris, 170 mM NaCl, 5% PEG. Scale bar = 10 µm.

D) Agarose gel results of Cytb rPCR (left) containing both dsDNA and ssDNA products and traditional PCR (right) containing only dsDNA product.

E) Radial distribution of ssDNA (red) and dsDNA (green) intensity in a representative four-component condensate (inset).

F) Schematic representation of the modeled macromolecules where protein TFAM and mtSSB are modeled as spheres and dsDNA and ssDNA as polymeric chains.

H) Strength of interactions for the four components in the minimalistic model.

### Supplementary Figure 5

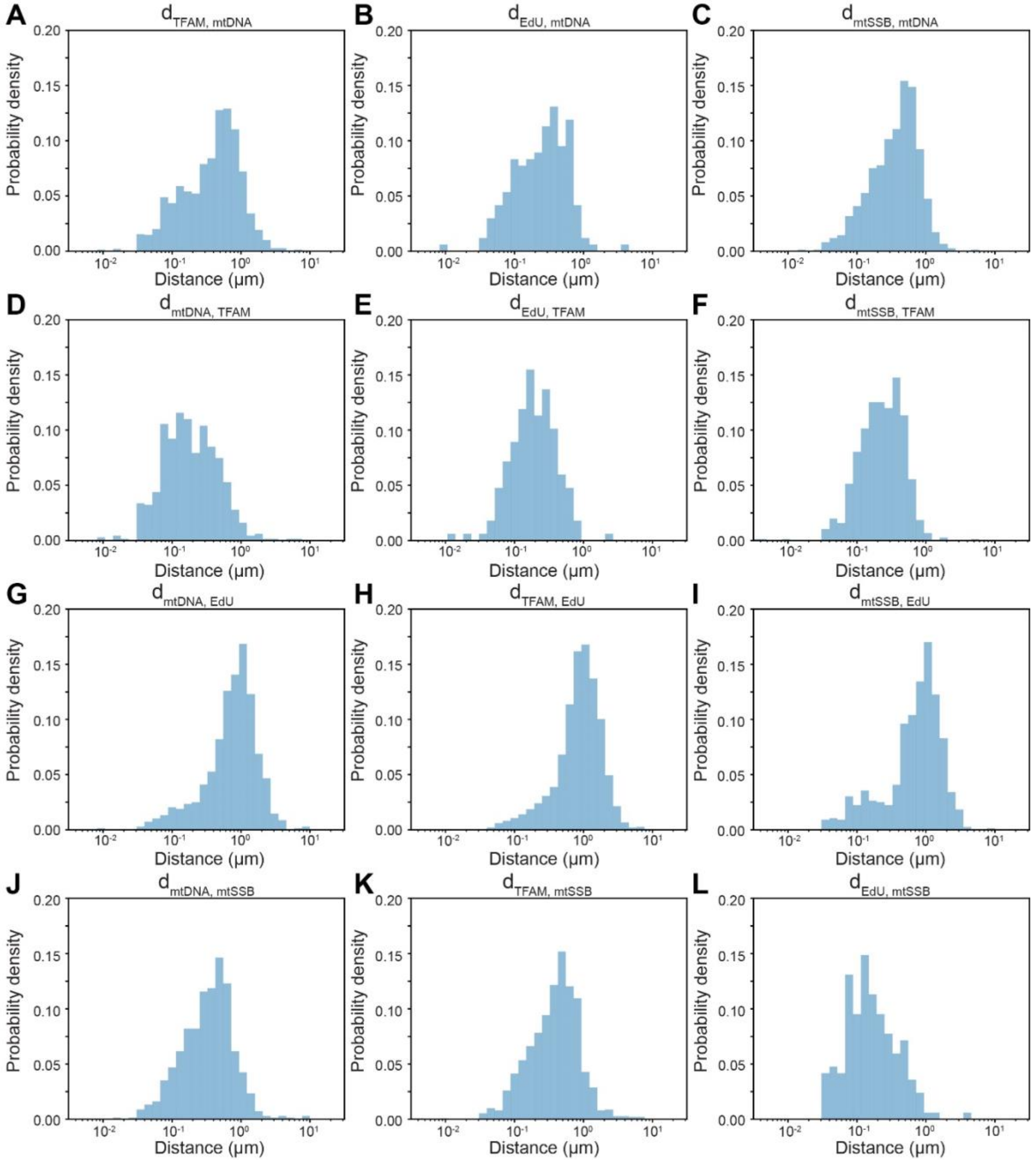


**Fig. S5 Probability density of nearest-neighbor analysis for mt-nucleoid components**

A-L) Probability density of nearest-neighbor distance, $d_{1,2}$, for mt-nucleoid components in HeLa cells labelled with anti-DNA, anti-mtSSB, EdU, and TFAM-mClover: $d_{TFAM,mtDNA}$ = nearest mtDNA punctum to a given TFAM punctum (A); $d_{EdU,mtDNA}$ = nearest mtDNA to EdU (B); $d_{mtSSB,mtDNA}$ = nearest mtDNA to mtSSB (C); $d_{mtDNA,TFAM}$ = nearest TFAM to mtDNA (D); $d_{EdU,TFAM}$ = nearest TFAM to EdU (E); $d_{mtSSB,TFAM}$ = nearest TFAM to mtSSB (F); $d_{mtDNA,EdU}$ = nearest EdU to mtDNA (G); $d_{TFAM,EdU}$ = nearest EdU to TFAM (H); $d_{mtSSB,EdU}$ = nearest EdU to mtSSB (I); $d_{mtDNA,mtSSB}$ = nearest mtSSB to mtDNA (J); $d_{TFAM,mtSSB}$ = nearest mtSSB to TFAM (K); $d_{EdU,mtSSB}$ nearest mtSSB to EdU (L).

**Supplementary Figure 6**

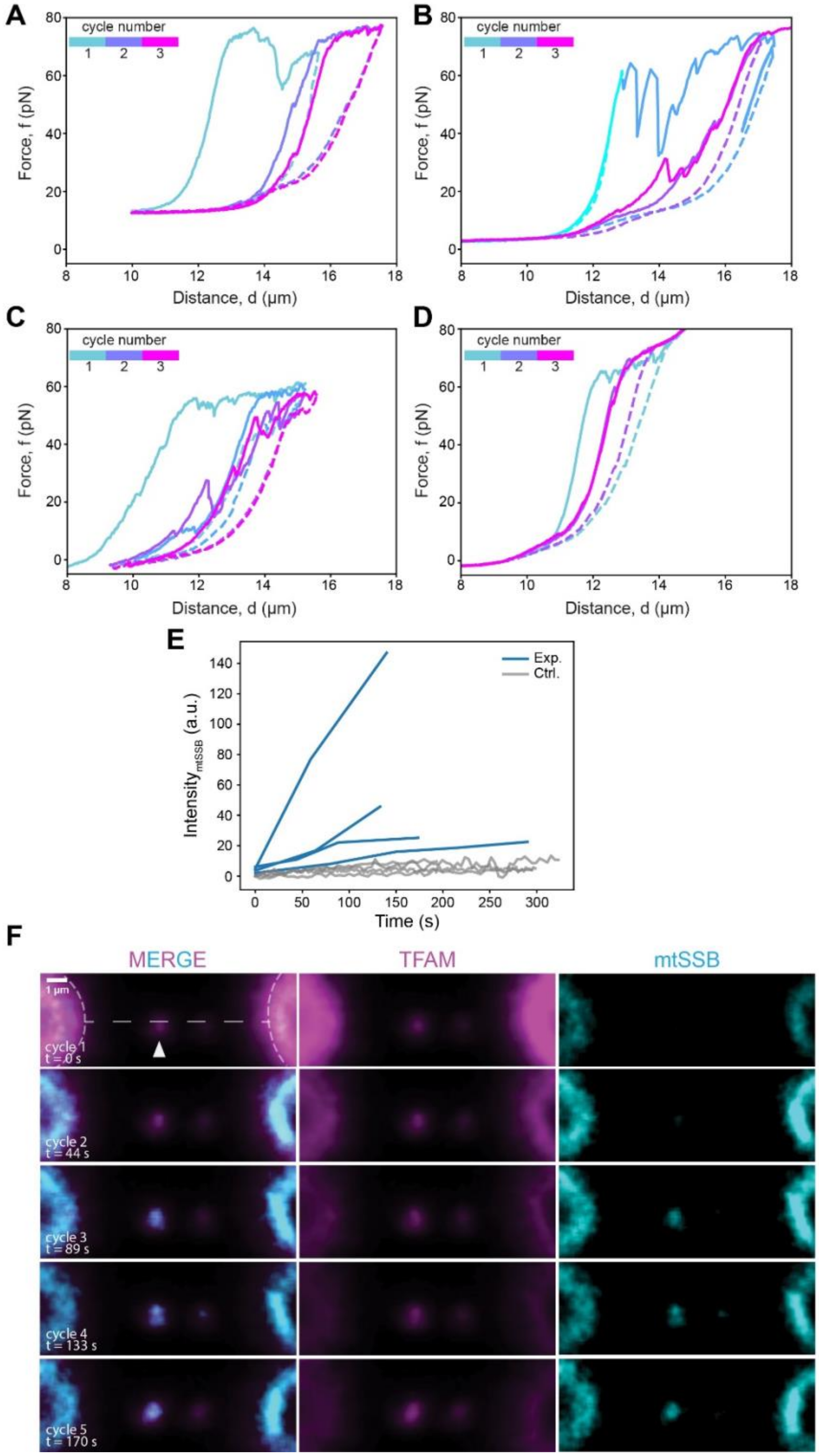

**Fig. S6 mtSSB partitioning into TFAM-DNA condensates upon repeated stretch-relax cycles**

A-D) Four representative force extension curves of mtSSB partitioning in TFAM-DNA condensates upon repeated stretch-relax cycles. λ-phage DNA was trapped between two beads and TFAM was applied to the flow cell channel. After condensate formation, DNA was moved to channel containing mtSSB in which multiple stretch/relax cycles were performed.

E) Change in mtSSB intensity in TFAM-DNA condensates over time. Blue lines: TFAM-bound DNA was moved to channel containing mtSSB and immediately underwent multiple stretch-relax cycles (experiment). Intensity of mtSSB in TFAM-DNA condensates was measured at the beginning of each cycle when end-to-end distance of DNA was roughly 8 μm. Gray lines: TFAM-bound DNA was moved to channel containing mtSSB and maintained at end-to-end distance ~8 μm for ~5 minutes without stretching (control). Intensity of mtSSB in TFAM-DNA condensates was measured for each frame.

F) Confocal image of mtSSB (cyan) partitioning in TFAM (magenta)-DNA condensates at different timepoints during pulling cycles. Scale bar = 1 μm.

**Supplementary Video Legends**

**Supplementary Video S1 Stretch-relax cycle of TFAM condensate bound to DNA.** DNA (green) was first immobilized between two beads trapped by an optical tweezer (C-trap, Lumicks). TFAM (magenta) was next applied to the channel and formed a condensate that compacted DNA. The optical trap was used to perform multiple stretch-relax cycles of the DNA and to measure resulting force. Scale bar = 1 μm.

**Supplementary Video S2 Partitioning of mtSSB into TFAM-DNA condensates upon repeated melting of DNA.** mtSSB (cyan) strongly partitioned into TFAM-DNA condensates (magenta) upon repeated stretch-relax cycles using optical tweezers (C-trap, Lumicks). Scale bar = 1 μm.

**Supplementary Video S3 Partitioning of mtSSB into TFAM-DNA condensates was reduced without force-induced DNA melting.** mtSSB (cyan) weakly partitioned into the TFAM-DNA condensate (magenta) as λ-DNA was held at end-to-end distance ~8 μm by optical tweezers (C-trap, Lumicks) without being stretched. Scale bar = 1 μm.
